# The MYCN/Aurora-A complex is a cyclin activating kinase for CDK12

**DOI:** 10.1101/2025.01.15.633177

**Authors:** Mareike Müller, Lorenz Eing, Leonie Uhl, Maximilian Schmitz, Iliyana Kaneva, Dominic P. Byrne, Daniel Fleischhauer, Selena G. Burgess, Sophia Fischer, Nick Gebauer, Vanessa Nancy-Portebois, Ines H. Kaltheuner, Stefanie Ahn Ha, Christina Schülein-Völk, Daniel Solvie, Mark W. Richards, Raphael Vidal, Matthias Brand, Dimitrios Papadopoulos, Richard Bayliss, Patrick A. Eyers, Claire E. Eyers, Matthias Geyer, Martin Eilers, Gabriele Büchel

## Abstract

Deregulated MYCN is a driver of aggressive pediatric and adult neuroendocrine tumors, but critical oncogenic processes downstream of MYCN remain poorly defined. In neuroblastoma, MYCN interacts with and activates the Aurora-A kinase. Here we show that Aurora-A is a CDK-activating kinase for CDK12 by phosphorylating T893 in the T-loop, thereby enhancing its kinase activity. Aurora-A-dependent activation of CDK12 controls phosphorylation of T4 of RNA polymerase and recruits transcription termination complexes, thereby preventing transcription-replication conflicts. Enhanced crosslinking and immunoprecipitation sequencing reveals that Aurora-A associates with splice sites on nascent RNA. RNA-bound Aurora-A is catalytically inactive. MYCN competes with RNA for binding to Aurora-A and displaces Aurora-A from RNA in cells, promoting its CDK12 kinase activity. Combining Aurora-A and CDK12 inhibition potently suppresses the growth of *MYCN*-amplified neuroblastoma cells and patient-derived xenografts. Our data demonstrate that an Aurora-A/CDK12-dependent transcription termination pathway is a critical and targetable dependency of MYCN-driven tumors.

## Introduction

Deregulated expression of each of three members of the MYC family of oncoproteins (MYC, MYCN, MYCL) typically occurs in distinct tumor entities, with MYCN being particularly prevalent in pediatric and adult neuroendocrine tumors ^1^. Even when expressed in tumors of the same entity, the different MYC paralogs drive subentities with distinct biological properties. A paradigmatic example is neuroblastoma, a pediatric tumor that arises from a highly proliferative population of neural crest cells ^2,3^. A subset of neuroblastomas is driven by an amplified *MYCN* oncogene and is characterized by rapid growth and frequent recurrence even after intensive chemotherapy ^4^. This leads to a poor prognosis for patients with *MYCN* amplified tumors and establishes an urgent need to identify the critical molecular processes underlying their aggressive clinical course ^4^.

Like MYC, MYCN forms a heterodimeric transcription factor with MAX, which binds broadly to active promoters and enhancers and affects the expression of a wide range of genes ^5,6^. While the effects of MYCN on individual genes are typically mild, they coalesce to generate a characteristic gene expression pattern in MYCN-driven tumors ^7,8^. The MYCN/MAX complex is prevalent in unperturbed cells. Upon stalling of RNA polymerase or perturbation of RNA processing, MYCN translocates onto nascent RNA, where it complexes with the nuclear exosome, a 3’-5’-RNA exonuclease complex, promoting degradation of nascent, non-polyadenylated RNA ^9^. RNA-bound MYCN complexes are prevalent during S phase and both RNA-binding of MYCN and exosomal RNA degradation prevent replication stress and DNA damage during S phase, suggesting that they coordinate transcription with DNA replication ^10^. During the S phase of the cell cycle, the presence of both the replication fork and RNA polymerases on the same template DNA can cause head-on and co-directional transcription-replication conflicts (TRCs) ^11,12^. Such conflicts pose a threat to genomic stability due to torsional stress that accumulates between the replication fork and RNA polymerases and because the stalling of replication and transcription induces the accumulation of aberrant, fragile nucleic acid structures, such as R-loops, resulting in double-strand breaks ^13^. Consequently, cells have evolved sophisticated mechanisms to resolve TRCs or prevent TRC-associated DNA damage ^12,14,15^. The mechanisms that maintain genomic stability during S phase are particularly important in tumor cells, which often proliferate under unfavorable conditions such as limiting nucleotide levels and rely on replication stress signaling to cope with the resulting problems ^16^. Since many drugs used in chemotherapy interfere with nucleotide metabolism and DNA replication, mechanisms that enhance resilience to stalling of replication forks or RNA polymerases may also contribute to chemotherapy resistance ^17^.

MYCN-driven tumors are particularly sensitive to depletion or inhibition of the Aurora-A serine-threonine kinase and Aurora-A inhibitors have high therapeutic efficacy in MYCN-driven tumor models ^18–20^. MYCN and Aurora-A interact directly, protecting MYCN from proteasomal degradation and stimulating the kinase activity of Aurora-A ^20,21^. Aurora-A phosphorylates many proteins during mitosis and is important for the proper assembly of the mitotic spindle ^22,23^. Surprisingly, depletion or inhibition of Aurora-A reveal that the kinase is critical for S phase progression and for protection from TRCs in MYCN-driven neuroblastoma cells ^5,24^. Combining inhibitors of Aurora-A and ATR, which stabilizes stalled replication forks ^12,14,15^, causes widespread and tumor-specific DNA damage in a model of MYCN-driven neuroblastoma, leading to massive apoptosis in tumors and a significant increase in the lifespan of tumor-bearing animals ^25^. The molecular mechanisms that enable Aurora-A to prevent TRCs, its substrates during S phase, and how Aurora-A function is affected by its interaction with MYCN are currently unknown. Here we combined the use of a specific Aurora-A inhibitor, LY-3295668 ^26,27^, with the expression of point mutants of Aurora-A that confer resistance to this inhibitor to identify specific substrates of Aurora-A. The subsequent analyses show that binding of MYCN activates Aurora-A to phosphorylate a pool of CDK12 that in turn phosphorylates RNAPII to promote transcription termination and prevent TRCs. Targeting this pathway with small molecules suppresses MYCN-, but not MYC-dependent neuroblastoma proliferation in culture and the growth of patient-derived xenografts in vivo, arguing that it provides a path to a molecularly targeted therapy of MYCN-driven tumors.

## Results

### Aurora-A phosphorylates the T-loop of CDK12

Treatment of *MYCN* amplified neuroblastoma cells with Aurora-A inhibitors or degraders strongly delays progression through S phase and results in S phase-specific DNA damage ^5,24^. While this argues that Aurora-A has specific phosphorylation targets that are critical for S phase progression, the identification of these targets has been difficult because many Aurora-A inhibitors also act on the closely related Aurora-B kinase ^28^. Here we used two strategies to overcome this problem. First, we used a highly specific Aurora-A inhibitor, LY-3295668 (hereafter abbreviated as LY668) ^26^. Immunoblots after immunoprecipitation (IP) of Aurora-A and Aurora-B showed that treatment of *MYCN* amplified IMR-5 neuroblastoma cells with LY668 abolished auto-phosphorylation of Aurora-A at T288 but did not reduce the auto-phosphorylation of Aurora-B at T232, in contrast to the Aurora-B specific inhibitor Barasertib (Figure 1A and Figure S1A). Second, we used two mutant alleles of murine Aurora-A, Aurora-A^T208D^ or Aurora-A^T208E^, that correspond to human Aurora-A^T217D^ and Aurora-A^T217E^. In human Aurora-A, these mutations confer enhanced resistance to several Aurora-A inhibitors ^29,30^ and we initially determined the effect of LY668 on recombinant murine Aurora-A kinase activity *in vitro*. Side-by-side comparison in the presence of ATP showed that both mutants have a significantly higher IC_50_ value (102.8 nM for Aurora-A^T208E^ and 220.0 nM for Aurora-A^T208D^ compared to 4.3 nM for Aurora-A^wt^, Figure 1B). We then established murine neuroblastoma NHO2A cell lines expressing either Aurora-A^wt^, Aurora-A^T208D^ or Aurora-A^T208E^ (Figure S1B). Consistent with an on-target effect of the inhibitor, treatment with LY668 suppressed the proliferation of NHO2A cells expressing Aurora-A^wt^ but had little effect on the growth of cells expressing Aurora-A^T208D^ and a weak effect on cells expressing Aurora-A^T208E^, in line with their relative IC_50_ values (Figure 1C, D). Since the IC_50_ value and the restoration of cell growth was significantly higher for Aurora-A^T208D^ than Aurora-A^T208E^ we used this mutant for the subsequent analyses. We confirmed that expression of Aurora-A^T208D^ conferred resistance to LY668 in two independent cell lines established from murine tumors, which developed in the TH-MYCN neuroblastoma *in vivo* model ^31^ (Figure S1C). To analyze the effects of LY668 on cell cycle progression, we performed BrdU/PI (bromodeoxyuridine/propidium iodide) flow cytometry experiments and found that inhibitor treatment induced the accumulation of both polyploid cells and of BrdU-negative cells in S phase, similar to the effects previously observed in *MYCN* amplified human neuroblastoma cells ^24^ (Figure S1D). Expression of the Aurora-A^T208D^ mutant, but not of Aurora-A^wt^, abrogated the cell cycle effects of LY668 treatment, resulting in a similar profile observed for the DMSO control. The cell cycle data, together with the *in vitro* and proliferation data validates that the cellular effects of LY668 are on-target. We therefore decided to perform global quantitative phospho-proteomic analyses to identify specific Aurora-A substrates.

**Figure 1:**
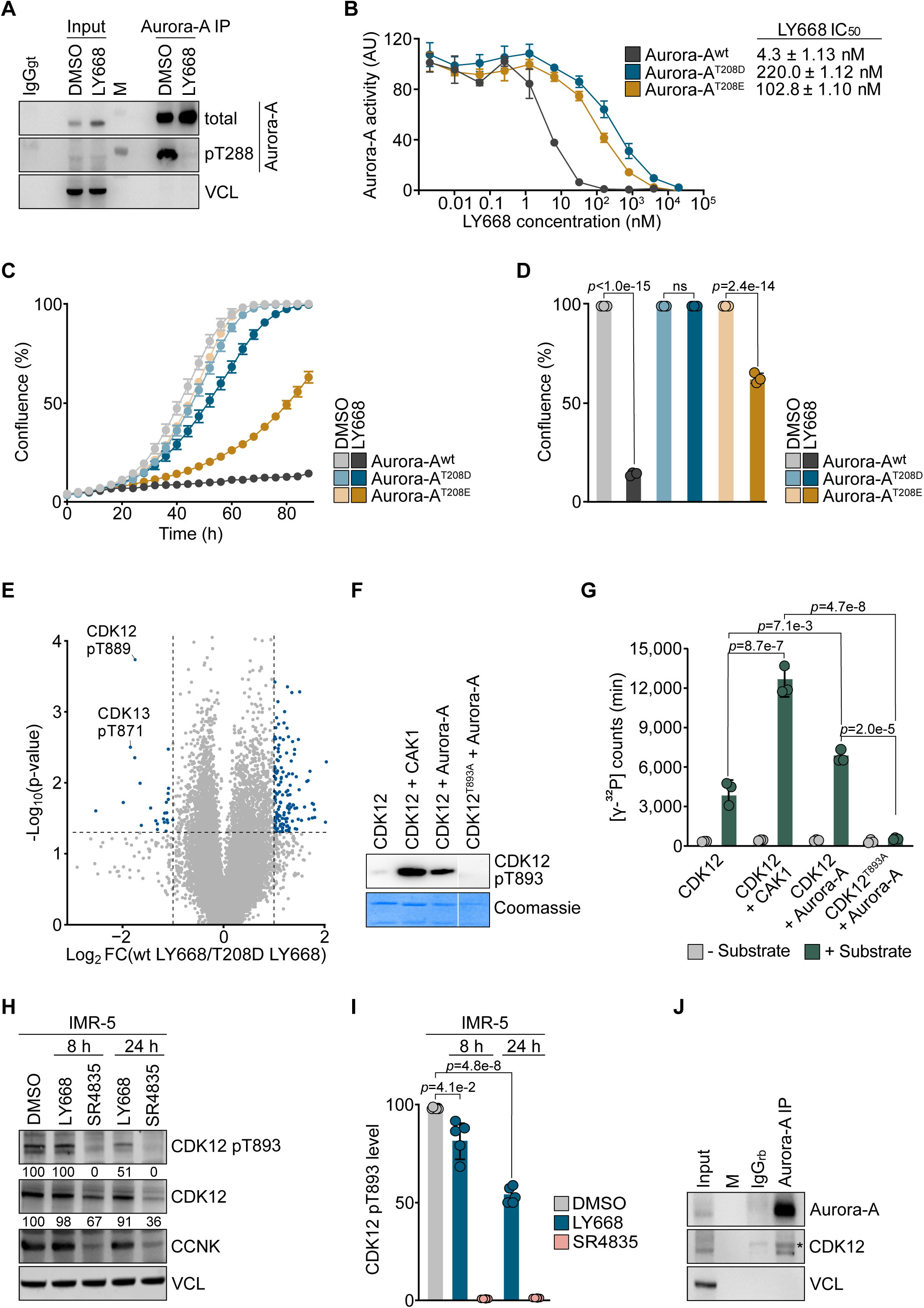
Aurora-A phosphorylates the T-loop of CDK12. **A.** Immunoblots of anti-Aurora-A IPs from IMR-5 cells treated with LY668 (1 µM, 24 h; those conditions were used unless otherwise specified). The input corresponds to 1% of the amount used for the precipitation. Non-specific IgG and VCL were used as controls (*n* = 3; *n* indicates the number of independent biological replicates, M = molecular weight marker). **B.** Graph shows *in vitro* kinase assay measuring murine Aurora-A activity in the presence of increasing concentrations of LY668. Purified full length Aurora-A^wt^, Aurora-A^T208D^ or Aurora-A^T208E^ were incubated with the fluorescently labelled peptide substrate in the presence of 1 mM ATP (*n* = 4). **C.** Growth curves of NHO2A cells expressing Aurora-A^wt^, Aurora-A^T208D^ or Aurora-A^T208E^ upon treatment with LY668. Data from three technical replicates of one representative experiment is shown as mean ± SD (*n* = 3). **D.** Endpoint confluence (t = 88 h) from growth curves of NHO2A cells described in panel C. Data shown as mean ± SD. *p* values calculated with one-way ANOVA (*n* = 3; ns, not significant). **E.** Volcano plot of quantitative phospho-proteomics data comparing the effect of LY668 on NHO2A cells expressing either Aurora-A^wt^ or Aurora-A^T208D^. The x axis displays phospho-peptide log_2_Fold change (FC) between cells expressing either Aurora-A^wt^ or Aurora-A^T208D^ following 8 h treatment with LY668. The y axis shows statistical significance (*n* = 3). **F.** *In vitro* CDK12 co-expression experiment with subsequent immunoblotting for CDK12 pT893. Coomassie is shown as a loading control (*n* = 1). **G.** *In vitro* kinase assay measuring CDK12 activity. Recombinant proteins were incubated with the substrate in presence of radiolabeled ATP. *p* values are calculated using a one-way ANOVA (*n* = 3). **H.** Immunoblots of indicated proteins in IMR-5 cells upon treatment with LY668 (1 µM) or SR4835 (50 nM) for 8 h or 24 h. VCL was used as a loading control. Quantification of CDK12 pT893 was normalized to total CDK12 level (*n* = 5). **I.** Quantification of CDK12 pT893 of biological replicates of immunoblots described in H. Data are shown as mean ± SD. *p* values are calculated using an unpaired two-sided t-test (*n* = 5). **J.** Immunoblots of anti-Aurora-A IPs from IMR-5 cells. The input corresponds to 1% of the amount used for the precipitation. Asterisks indicate unspecific bands. Non-specific IgG and VCL were used as controls (*n* = 3, M = molecular weight marker). See also Figure S1.

Comparing inhibitor-treated NHO2A cells that express Aurora-A^wt^ with treated cells expressing Aurora-A^T208D^ identified 24 significantly affected phosphorylation sites (log_2_FC<-1, *p* value<0.05, Table S1). Of those, T889 in murine CDK12 (cyclin-dependent kinase 12) and T871 in CDK13 showed the most significant decrease in phosphorylation in cells expressing Aurora-A^wt^ (Figure 1E). Controls established that phosphorylation at either site did not differ significantly between cells expressing Aurora-A^wt^ or Aurora-A^T208D^ in the absence of LY668 (Figure S1E). Surprisingly, we also found a group of phosphorylation sites that were increased in LY668-treated cells expressing Aurora-A^wt^. Annotation of these sites showed that almost all are canonical mitotic targets that are shared between Aurora-A and Aurora-B (Figure S1F). As described before, treatment of cells expressing Aurora-A^wt^, but not Aurora-A^T208D^ with LY668 resulted in the accumulation of cells in the S/G2 phase, when Aurora-B is active (Figure S1D). As LY668 does not inhibit Aurora-B (Figure S1A), this suggested that these mitotic targets are phosphorylated by Aurora-B in LY668-treated cells. Supporting this hypothesis, we found that LY668 increased Aurora-B autophosphorylation at T232 and increased H3S10 phosphorylation, a well-established substrate of both Aurora-A and Aurora-B (Figure S1A, G). In contrast, treatment with the Aurora-B inhibitor Barasertib abolished Aurora-B phosphorylation at T232 (Figure S1A) and co-treatment abolished histone H3S10 phosphorylation in LY668-treated cells (Figure S1G). We concluded that T889 in CDK12 and T871 in CDK13 are specific targets of Aurora-A in neuroblastoma cells.

T889 in CDK12 and T871 in CDK13 are located within the T-loop, which is targeted by cyclin-activating kinase (CAK) ^32^. Importantly, T-loop phosphorylation by CAK is essential for cyclin binding and catalytic activity ^33^. CDK12 and CDK13 are structurally very similar, and we focused on CDK12 in the subsequent analyses ^34,35^. To test whether Aurora-A can directly phosphorylate T889, we incubated recombinant active human CDK12, which was expressed together with cyclin K, with Aurora-A *in vitro* and probed the reactions with a phospho-specific antibody that recognizes human pT893 CDK12, which is homologous to T889 in murine CDK12 (Figure S1H). Western blot analysis confirmed that Aurora-A is able to phosphorylate T893 *in vitro* with a slightly lower efficiency than recombinant CAK1, the canonical kinase that phosphorylates the T-loop in several CDKs ^36^ (Figure 1F). A point mutant, CDK12^T893A^, was used to validate the specificity of the antibody. Furthermore, *in vitro* kinase assays using a CTD peptide of RNAPII as substrate showed that incubation with Aurora-A enhanced the catalytic activity of CDK12, but not of CDK12^T893A^, relative to control (Figure 1G).

Next, we used the phospho-specific antibody that recognizes pT893 in human CDK12 to determine the phosphorylation status in neuroblastoma cells, which showed that inhibition of Aurora-A reduced phosphorylation at T893 of CDK12 in human IMR-5 cells (Figure 1H, I). Similarly, pT889 in murine NHO2A cells expressing Aurora-A^wt^ was reduced, while T889 phosphorylation was unaffected in NHO2A cells expressing Aurora-A^T208D^, confirming that the effect of the inhibitor on pT889 is on-target (Figure S1I, J). Notably, inhibition of Aurora-A did not affect the entire pool of cellular CDK12 in either human or mouse cells. Rather, phosphorylation of CDK12 at the T-loop was decreased by 40-50% in both cell types, even though Aurora-A was essentially inactive after LY668 treatment (as monitored by T288 autophosphorylation) (Figure 1A). In contrast, treatment of cells with the CDK12 inhibitor SR4835 completely abolished phosphorylation at T893 and led to a significant decrease in the levels of CDK12 itself and cyclin K (CCNK), the cognate cyclin of CDK12, consistent with previous observations that SR4835 can act as a molecular glue for CDK12 ^37,38^. In line with the ability of Aurora-A to phosphorylate CDK12, we found that CDK12 was present in endogenous Aurora-A IPs (Figure 1J). Taken together, the data show that Aurora-A phosphorylates approximately 40-50% of the cellular CDK12 pool at T893 identifying Aurora-A as a CAK of CDK12.

### Phosphorylation of T4 in the CTD of RNAPII by CDK12 depends on Aurora-A

CDK12 enhances the expression of genes expressed during the S phase by suppressing intronic polyadenylation ^39^ and promotes the splicing of proximal introns ^40^. To determine whether both processes are downstream of Aurora-A in neuroblastoma cells, we performed RNA sequencing in IMR-5 cells treated with the CDK12 inhibitor SR4835 or the Aurora-A inhibitor LY668. While inhibition of CDK12 by SR4835 had widespread effects on gene expression, treatment with LY668 elicited only minor changes (Figure S2A). To analyze the role of Aurora-A in the expression of DNA damage response genes and intronic polyadenylation ^34,39^ we evaluated a panel of genes using quantitative PCR following reverse transcription (RT-qPCR). Since CDK12 suppresses the use of intronic polyadenylation during S phase ^39^, we used two sets of primer pairs. The first set is located after the 3’-outermost exon of a gene, reflecting the distal polyadenylation site (PA). The second set of primers probes the use of intron-located PA sites. While the isoforms of genes containing the distal PA sites was strongly downregulated in SR4835 treated cells, LY668 treatment showed only a slight reduction (Figure S2B). We then checked the usage of intronic PA sites and found that expression of isoforms that use intronic PA sites increases after treatment with SR4835, but not after Aurora-A inhibition (Figure S2C). To analyze global changes in splicing, we performed an rMATS analysis ^41,42^. This showed that direct inhibition of CDK12 using SR4835 caused significant increases in aberrant splicingm which is shown here for intron retention, while Aurora-A inhibition with LY668 had very little effects on the splicing events that were analyzed (Figure S2D). We concluded that transcription of DNA damage genes, suppression of intronic polyadenylation and splicing of proximal introns are mediated by a pool of CDK12 that does not depend on Aurora-A-dependent activation.

CDK12 directly phosphorylates the carboxy-terminal domain (CTD) of RNAPII ^43^. To better understand the role of Aurora-A dependent phosphorylation of CDK12, we performed spike-normalized chromatin-immunoprecipitation and sequencing (ChIP-Rx) of total RNAPII as well as RNAPII phosphorylated on the CTD at S2 or T4, which are both marks of elongating RNAPII. Inhibitor treatment had minor effects on the levels of RNAPII (Figure S3A). Inspection of individual promoters showed that chromatin association of total RNAPII decreased upon SR4835, while inhibition of Aurora-A had only a minor effect (Figure 2A and Figure S3B). Genome-wide analysis of the elongating form of RNAPII (RNAPII pS2) revealed a significant decrease after treatment with SR4835 while LY668 resulted only in a minor reduction (Figure 2A, B and Figure S3B). In sharp contrast, RNAPII pT4 occupancy was significantly downregulated after both SR4835 and LY668 treatment (Figure 2A, B, and Figure S3B). RNAPII T4 phosphorylation progressively increases during elongation and has been linked to 3’-end processing of RNA and transcription termination ^44–46^. To confirm this observation, we tested the enrichment of RNAPII pT4 by ChIP-qPCR with different primers covering different locations of the *KPNB1* locus upon inhibitor treatment. We observed that RNAPII pT4 increases throughout the gene (Figure S3C) and that occupancy at the distal end of several genes is strongly decreased upon LY668 treatment (Figure 2C and Figure S3C). To confirm that this effect is mediated by Aurora-A, we used the established murine neuroblastoma cell lines NHO2A expressing either Aurora-A^wt^ or Aurora-A^T208D^ and performed ChIP-qPCR upon LY668 treatment. While RNAPII pT4 decreased at the transcription end site (TES) upon inhibitor treatment of cells expressing Aurora-A^wt^, pT4 was unaffected in cells expressing Aurora-A^T208D^ (Figure 2D). To evaluate if RNAPII T4 is a direct target of CDK12 phosphorylation we performed *in vitro* kinase assays. This confirmed that the CDK12/CCNK complex induces T4 phosphorylation of RNAPII and that this is abolished upon treatment with SR4835 (Figure 2E). Importantly, Aurora-A by itself did not phosphorylate T4 in *in vitro* kinase assays (Figure S3D). We conclude that Aurora-A is required for CDK12-dependent phosphorylation of RNAPII T4, but not S2. To explore possible consequences for 3’-end processing of RNA and transcription termination, we analyzed the proximity of RNAPII to different termination complexes using proximity ligation assays (PLAs). We analyzed the proximity of RNAPII to PCF11 and CPSF73, which are components of the canonical polyadenylation and termination complexes ^47^, to ZC3H4, which helps terminate unproductive termination ^48^, and to INTS11, a component of the integrator complex that targets paused RNAPII for termination ^49^. Intriguingly, these experiments showed that both Aurora-A and CDK12 inhibition attenuated the proximity of all termination complexes to RNAPII. We concluded that Aurora-A activates a pool of CDK12 that enables RNAPII to engage different termination complexes (Figure 2F, G and Figure S3E).

**Figure 2:**
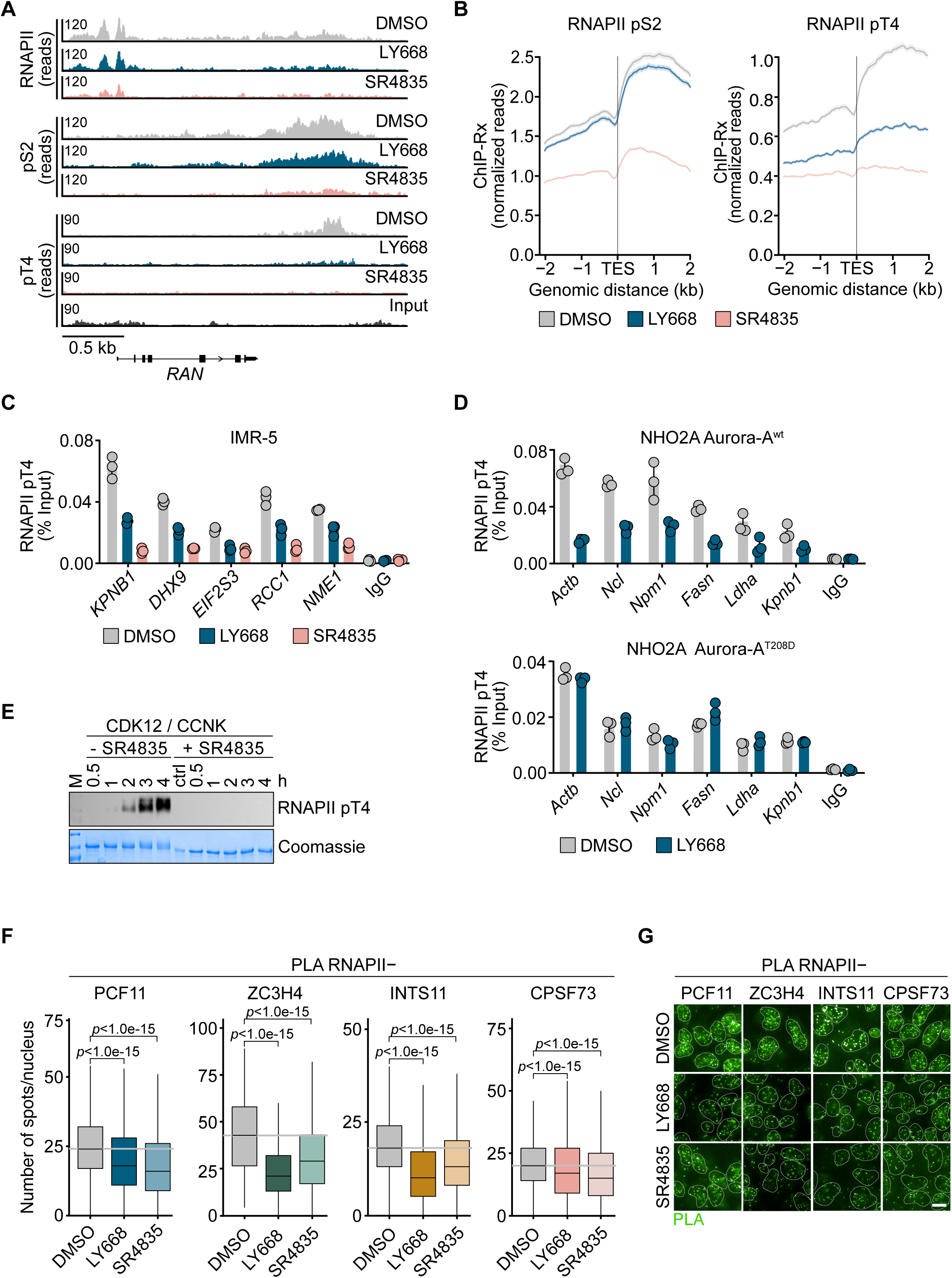
Phosphorylation of RNAPII T4 by CDK12 is dependent on Aurora-A. **A.** Genome browser tracks of normalized reads at the *RAN* locus showing chromatin association of RNAPII (total, pS2 or pT4) in IMR-5 cells upon treatment with LY668 (1 µM), SR4835 (50 nM) for 24 h or DMSO as a control (*n* = 2 for total RNAPII and pS2; *n* = 1 for pT4). **B.** Global average read density of RNAPII pS2 or pT4 ChIP-Rx centered around the transcription end site (TES) of all expressed genes as described in A. Data show mean ± SEM (N = 14,704 genes analyzed, *n* = 2 for pS2; *n* = 1 for pT4). **C.** RNAPII pT4 ChIP-qPCR at indicated loci in IMR-5 cells after treatment as indicated for 24 h. IgG was used as negative control. Shown are technical triplicates with the mean of one representative experiment (*n* = 3). **D.** RNAPII pT4 ChIP-qPCR at indicated loci in NHO2A cells expressing Aurora-A^wt^ (top) or Aurora-A^T208D^ (bottom) after treatment with LY668 for 8 h. IgG was used as negative control. Shown are technical triplicates with the mean of one representative experiment (*n* = 3). **E.** *In vitro* kinase activity experiment with subsequent immunoblotting for RNAPII-CTD activity (pT4) in absence or presence of SR4835 at stated time points. Coomassie staining is shown as loading control (*n* = 1). **F.** Boxplot of single-cell analysis of nuclear proximity ligation assay (PLA) foci between RNAPII and PCF11, ZC3H4, INTS11, or CPSF73 in IMR-5 cells of one representative PLA experiment. *p* values were calculated comparing the PLA signal of all analyzed cells using an Wilcoxon rank sum test compared to the control condition (*n* = 3 for PCF11, ZC3H4, *n* = 2 for INTS11, *n* = 1 for CPSF73). **G.** Representative PLA images from the experiment quantified in panel F showing PLA foci after treatment with LY668, SR4835 or DMSO as control. Scale bar indicates 10 µm. See also Figure S2 and S3.

### Inhibition of Aurora-A or CDK12 rapidly induces transcription-replication conflicts

We next set out to explore whether regulation of T4 phosphorylation and recruitment of termination complexes to RNAPII have a function in cell cycle progression of neuroblastoma cells. Earlier studies using MLN8237 (Alisertib) or an MLN8237-based Aurora-A-PROTAC indicated that Aurora-A is critical for S phase progression in MYCN-driven neuroblastoma cells and protects them from TRCs ^5,24,25^. To confirm whether this is observed using LY668 and determine whether it also depends on CDK12, we initially tested the effect of LY668 or SR4835 on cell cycle progression in IMR-5 cells. BrdU/PI flow cytometry analysis showed that inhibition of both Aurora-A and CDK12 strongly perturbed DNA synthesis and led to the accumulation of BrdU-negative cells with a DNA content between 2n and 4n, demonstrating that they did not synthesize DNA and therefore have not duplicated the entire genome (Figure 3A, B and Figure S4A). Very similar effects were observed using quantitative immunofluorescence of EdU-labelled cells (Figure S4B), confirming that both Aurora-A and CDK12 are required for S phase progression.

**Figure 3:**
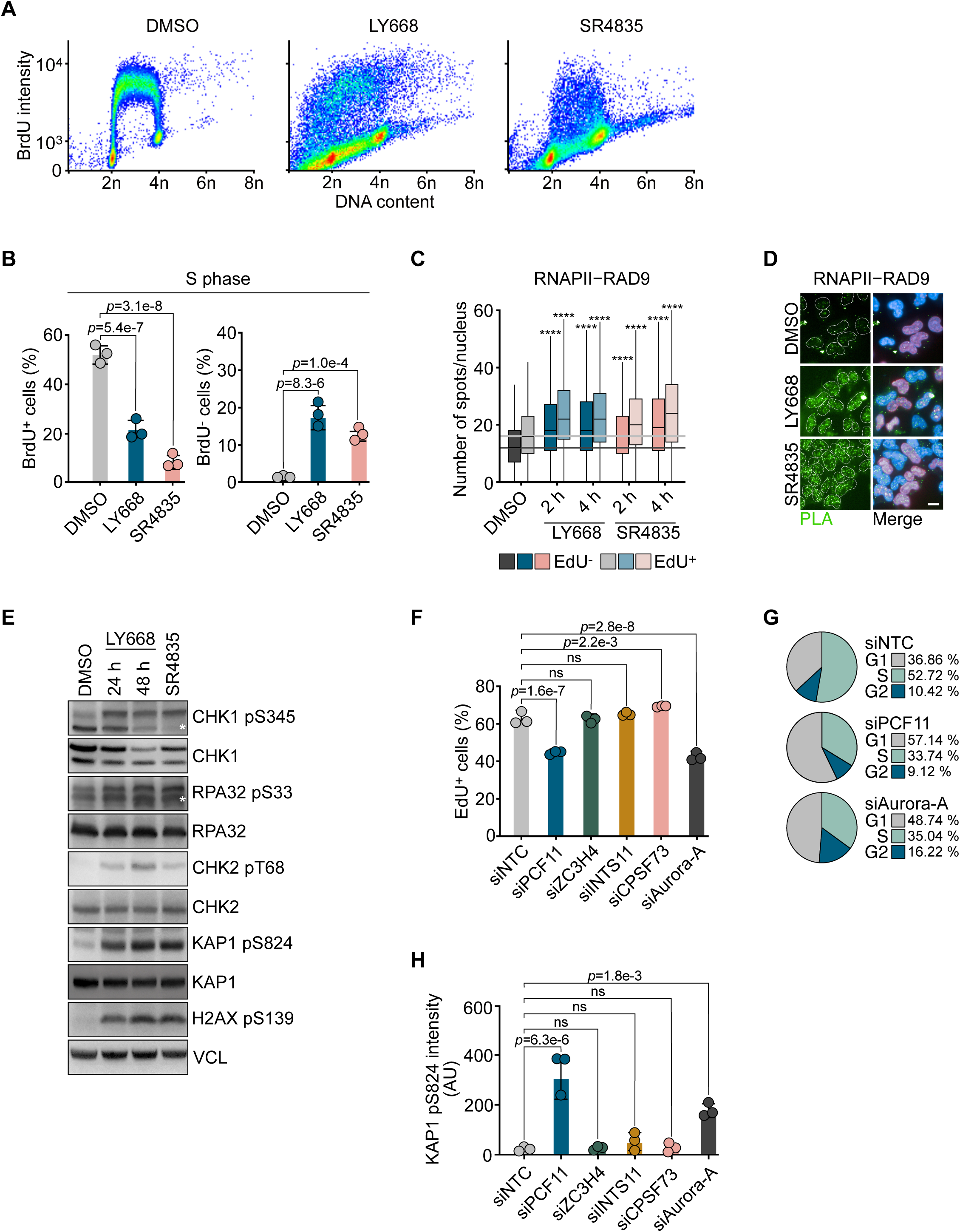
Inhibition of Aurora-A and CDK12 induce transcription-replication conflicts. **A.** BrdU/PI flow cytometry profiles of IMR-5 cells treated with LY668 (1 µM), SR4835 (50 nM) for 24 h or DMSO as a control (*n* = 3). **B.** Quantification of BrdU^+^ (left) or BrdU^-^ (right) cell populations of FACS analysis described in A. Data are shown as mean ± SD. *p* values were calculated using an one-way ANOVA (*n* = 3). **C.** Boxplot of single-cell analysis of nuclear PLA foci between RNAPII and RAD9 in IMR-5 cells treated with LY668 (1 µM) or SR4835 (50 nM) for 2 h or 4 h or DMSO as control. Conditions are stratified for EdU incorporation. *p* values were calculated comparing the PLA signal of all analyzed cells using an Wilcoxon rank sum test compared to corresponding control condition (\*\*\*\**p* value <1.0e-15, *n* = 3). **D.** Representative images showing PLA foci (left) and PLA, EdU and Hoechst signal as merge (right) of cells quantified in panel C. Scale bar indicates 10 µm (*n* = 3). **E.** Immunoblots of indicated proteins in IMR-5 cells upon treatment with LY668 (1 µM, 24 h or 48 h) or SR4835 (50 nM, 24 h). Asterisks indicate unspecific bands. VCL was used as a loading control (*n* = 3). **F.** Analysis of EdU^+^ cells upon transfection with indicated siRNAs in IMR-5 cells. *p* values were calculated using an one-way ANOVA (*n* = 3). **G.** Representative cell cycle distribution of IMR-5 cells determined with immunofluorescence upon transfection with siPCF11, siAurora-A or siNTC as control (*n* = 3). **H.** Intensity of KAP1 pS824 in IMR-5 cells upon transfection with indicated siRNAs. *p* values were calculated using an one-way ANOVA (*n* = 3). See also Figure S4.

To determine whether the dependence on Aurora-A for S phase progression correlates with the occurrence of TRCs in inhibitor-treated cells, we performed PLAs between RNAPII and RAD9, which is a marker for stalling replication forks ^50^. These assays showed that treatment of IMR-5 cells with LY668 or SR4835 caused an accumulation of stalled replication forks close to RNAPII within 2 hours of inhibitor treatment, with the effect being more prominent in EdU-positive cells (Figure 3C, D and Figure S4C). To explore the downstream consequences of these events, we analyzed several DNA damage markers in IMR-5 cells. Immunoblots using phospho-specific antibodies showed that inhibition of Aurora-A or CDK12 induced phosphorylation of CHK1 (pS345) and RPA32 (pS33), which are phosphorylated by the ATR kinase that is activated in response to stalling of replication forks, and of KAP1 (pS824), CHK2 (pT68) and H2AX (pS139), which are predominantly phosphorylated by the ATM kinase or DNA-PK that monitor double-strand breaks (Figure 3E). Surveying a panel of neuroblastoma cell lines revealed that the DNA damage responses induced by inhibition of either Aurora-A or CDK12 were more pronounced in *MYCN* amplified neuroblastoma cells than in *MYCN* non-amplified cells, arguing for a critical role of MYCN in this pathway (Figure S4D, E). Finally, we asked whether any of the termination complexes that are recruited to RNAPII in an Aurora-A dependent manner is required for suppressing DNA damage in S phase. To do so, we depleted each of these factors using siRNA (Figure S4F) and analyzed the impact on S phase progression using EdU labeling. These assays revealed that depletion of PCF11 mirrors the defects in EdU incorporation of Aurora-A depletion, whereas downregulation of other termination factors had no negative impact on EdU incorporation (Figure 3F, G and Figure S4G). We subsequently analyzed DNA damage by immunofluorescence staining of pKAP1 (Figure 3H). Depletion of PCF11 by siRNA also induced an increase in KAP1 phosphorylation similar to that observed upon Aurora-A depletion. PCF11 is a component of the canonical polyadenylation and termination complex ^47^ but also accumulates in promoter-proximal regions when RNAPII is paused ^51^, suggesting that it also participates in other termination pathways. Collectively, these data show that Aurora-A and CDK12 are required to prevent TRCs in *MYCN* amplified neuroblastoma and that this correlates with their ability to promote transcription termination.

### Aurora-A binds to splice sites on nascent RNA

We hypothesized that Aurora-A is likely to be spatially close to RNAPII under conditions where its activity on CDK12 is relevant and therefore used PLAs to analyze the proximity of Aurora-A to RNAPII under stress- or damage-inducing conditions (Figure 4A, B and Figure S5A). These assays revealed a robust PLA signal in unperturbed IMR-5 cells, which was enhanced in EdU-positive cells (Figure 4A and Figure S5A). Incubation of cells with compounds that delay S phase progression or block DNA replication, such as hydroxyurea (HU), aphidicolin (APH) or thymidine (THD) block, led to a significant increase in the PLA signals (Figure 4A, B). In addition, inhibition of topoisomerase I by irinotecan and of the FACT histone chaperone complex by CBL0137 ^52^ also increased the proximity between RNAPII and Aurora-A (Figure S5A). The effects of all inhibitors were more pronounced in EdU-positive cells (Figure S5A). Since irinotecan and CBL0137 inhibit transcriptional elongation, these data suggest that a delay in both DNA replication and in transcription elongation, resulting in a slowing of RNAPII, enhance the proximity of Aurora-A to RNAPII.

**Figure 4:**
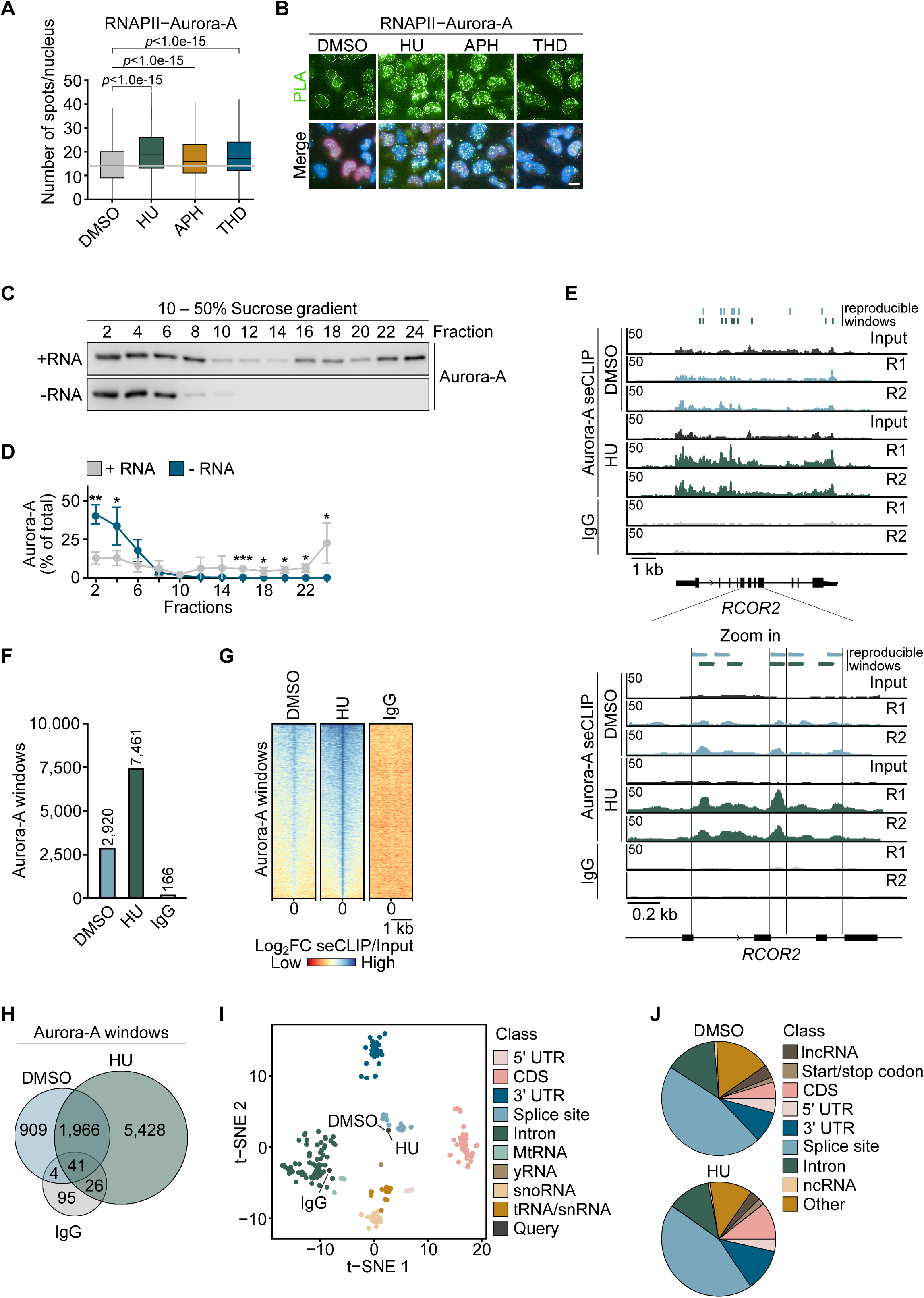
Aurora-A binds to nascent RNA. **A.** Boxplot of single-cell analysis of nuclear PLA foci between RNAPII and Aurora-A in IMR-5 cells treated with Hydroxyurea (HU, 3.5 mM, 2 h), Aphidicolin (APH, 1 µM, 4 h), or thymidine (THD, 2 mM, 10 h). *p* values were calculated by comparing the PLA signal of all analyzed cells with the corresponding control using an Wilcoxon rank sum test (*n* = 2). **B.** Representative PLA images showing PLA signal and merge with EdU and Hoechst in cells described in A. Scale bar indicates 10 µm (*n* = 2). **C.** Immunoblot of Aurora-A sedimentation patterns on 10-50% sucrose gradients from IMR-5 cells with (top) and without (bottom) RNA (*n* = 3). **D.** Quantification of Aurora-A sedimentation patterns described in C. Data are shown as mean ± SD. *p* values were calculated using an unpaired two-sided t-test (\**p* value < 0.05; \*\**p* value<0.01; \*\*\**p* value<0.001). **E.** (Top) Genome browser tracks of Aurora-A seCLIPs at the *RCOR2* locus. IMR-5 cells were treated with HU (3.5 mM, 2 h) or DMSO as a control. Input and IgG are shown as control. Bars on top show reproducible enriched windows. (Bottom) Zoom in into the *RCOR2* locus (*n* = 2, R = replicate). **F.** Bar graph depicting numbers of reproducible enriched windows of Aurora-A seCLIPs of replicate conditions described in E. Windows in IgG seCLIP are shown as control. **G.** Heatmaps showing the log_2_FC of Aurora-A seCLIPs over the respective size-matched input in depicted conditions. **H.** Venn diagram of reproducible windows of merged replicates of Aurora-A seCLIPs for HU treated and DMSO treated IMR-5 cells and IgG seCLIP as control. **I.** t-distributed stochastic neighbor embedding (t-SNE) analysis comparing Aurora-A binding to publicly available (ENCODE) eCLIP data of RNA-binding proteins. IgG seCLIP was performed as control. **J.** Pie chart showing enriched windows for RNA species classes for stated Aurora-A seCLIP conditions. See also Figure S5.

Since we recently found that the proximity of MYCN to RNAPII is mediated by its association with nascent RNA ^9^, we speculated that Aurora-A may similarly interact with nascent RNA during transcription termination. We initially performed sucrose gradient centrifugation of cell lysates in control conditions and upon treatment with a mixture of different RNases to degrade RNA. This showed that Aurora-A, like MYCN, forms high molecular weight complexes in an RNA-dependent manner (Figure 4C, D). Next, we performed single-end enhanced cross-linking and immunoprecipitation (seCLIP) experiments, in which proteins are crosslinked to RNA by UV irradiation and bound RNA molecules are subsequently recovered by immunoprecipitation and sequenced ^53^. These experiments revealed a robust signal for Aurora-A on many transcribed genes (Figure 4E and Figure S5B). There was no significant enrichment of specific RNAs in seCLIP experiments using a control IgG, confirming the specificity of the signal (Figure 4E-H and Figure S5B-D). Bioinformatic analyses identified 2,920 reproducible binding sites (“windows”) of Aurora-A on RNA in untreated cells (Figure 4F, H and Figure S5D). Arresting cells in S phase using HU increased the number of Aurora-A RNA binding sites to 7,461, with a large overlap with the binding sites in control cells (Figure 4F, H and Figure S5D). Comparing Aurora-A binding to known RNA binding proteins (RBPs) revealed that sequences bound by Aurora-A in both control and HU-treated cells cluster with multiple RBPs that associate with splice sites (Figure 4I). Similarly, annotation of data peaks based on their location showed that over 50% of Aurora-A binding sites were localized at or proximal to splice sites of protein-coding genes (Figure 4J and Figure S5E). This was confirmed by inspection of individual genes (Figure 4E, bottom). We concluded that Aurora-A robustly binds to RNA with a preference for splice sites and that RNA binding is enhanced in S phase-arrested cells.

### MYCN displaces Aurora-A from RNA and activates Aurora-A kinase activity

Aurora-A forms a complex with MYCN during S phase ^5^, and MYCN translocates from its cognate binding sites at promoters onto nascent RNA when RNAPII stalls and intronic RNA accumulates ^9^. MYCN binds RNA through MYCBoxI ^9^, the same domain involved in the MYCN-Aurora-A interaction ^21^, suggesting that complex formation of Aurora-A and MYCN may affect RNA binding by either protein. To test whether Aurora-A directly binds RNA, we incubated recombinant purified Aurora-A (aa 122-403) as well as a covalently cross-linked protein comprising of this domain of Aurora-A and a MYCN peptide (aa 28-89) which encompasses both the Aurora-A and RNA binding regions of MYCN (Aurora-A:MYCN cross-link ^54^) with a fluorescently labeled UG-rich RNA oligonucleotide that was previously shown to bind MYCN (“UG^high^”) ^9^ and evaluated concentration-dependent interactions by gel mobility shift assays. We have shown previously that the structure of the cross-linked Aurora-A:MYCN complex is identical to that of the native Aurora-A/MYCN complex ^54^. We observed that Aurora-A interacts with RNA and that cross-linking of Aurora-A to a MYCN peptide reduced its RNA affinity, suggesting that MYCN antagonizes Aurora-A binding to RNA (Figure 5A, B and Figure S6A). To further investigate the role of MYCN in RNA binding of Aurora-A, we performed *in vitro* co-precipitation and competition assays, in which we added increasing amounts of MYCN to Aurora-A (aa 1-403) and monitored Aurora-A’s ability to bind UG^high^ RNA oligonucleotide. In these assays, Aurora-A was recovered on beads carrying RNA, but not on control beads. Addition of MYCN abrogated RNA binding of Aurora-A (Figure 5C) and titration showed that equimolar amounts of MYCN were sufficient to inhibit RNA binding (Figure 5D). Notably, an ATP-competitive inhibitor of Aurora-A (SCH1473759 ^55^) had no effect on the ability of the kinase to bind RNA, suggesting that RNA binding and its inhibition by MYCN are not due to occupancy and accessibility of the ATP-binding pocket of Aurora-A (Figure 5C).

**Figure 5:**
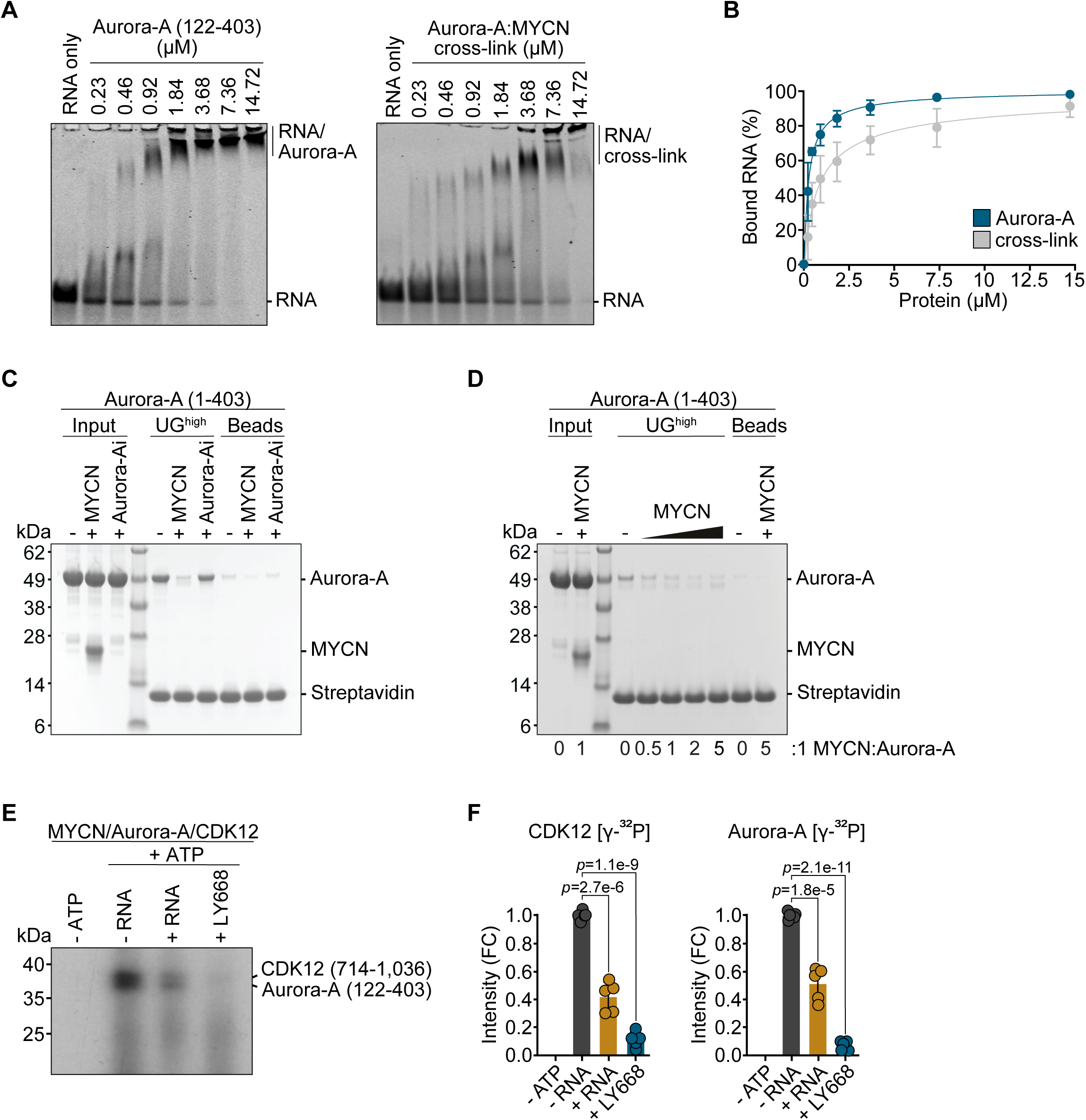
MYCN-Aurora-A complex binds RNA less efficiently. **A.** Electrophoretic mobility shift assay (EMSA) using Aurora-A (left) or Aurora-A:MYCN cross-link (right) incubated with 5 nM UG^high^ RNA. The RNA probe was visualized by its Cy5 label (*n* = 5). **B.** Quantification of bound RNA in replicates of EMSAs described in A. Data are shown as mean ± SD (*n* = 5). **C.** SDS-PAGE of *in vitro* co-precipitation assays reporting binding interactions between biotinylated UG^high^ RNA oligonucleotide and proteins as indicated. UG^high^ RNA was incubated with the proteins before capture on Streptavidin beads. SCH1473759, a type 1 competitive Aurora-A inhibitor (Aurora-Ai), was added at a 1:1 kinase:drug stoichiometry (*n* = 3). **D.** SDS-PAGE analysis of *in vitro* co-precipitation assays to observe binding of unphosphorylated Aurora-A (1-403) protein to UG^high^ RNA oligonucleotide in the presence of increasing concentrations of MYCN (1-137) protein (*n* = 3). **E.** *In vitro* kinase assay measuring Aurora-A and CDK12 activity. Recombinant proteins were incubated in presence of radiolabeled ATP. First lane shows background signal without addition of ATP (*n* = 5). **F.** Quantification of kinase assay described in E. Data show mean ± SEM. *p* values were calculated using an unpaired two-sided t-test (*n* = 5). See also Figure S6.

To be a fully, active kinase, Aurora-A requires interaction with a co-activator. In mitosis, this is TPX2 ^56,57^ but MYCN also activates Aurora-A *in vitro* ^21^ and we confirmed this observation using a luminescence-based kinase assay (Figure S6B). To explore how binding of Aurora-A to RNA affects its activity towards CDK12, we incubated Aurora-A and MYCN with recombinant CDK12 in the presence and absence of the UG^high^ RNA oligonucleotide. In these assays, we observed both Aurora-A autophosphorylation and phosphorylation of CDK12 (Figure 5E and Figure S6C). Upon addition of RNA, phosphorylation of both targets significantly decreased, indicating that Aurora-A activity is inhibited by binding to RNA. As a control for specificity, we inhibited Aurora-A activity using LY668, which abolished the phosphorylation signal (Figure 5E, F).

These data suggest that RNA-bound Aurora-A is inactive and that MYCN displaces Aurora-A from RNA in cells. In human neuroblastoma cells, a mutation in MYCN that replaces three positively charged residues in MYCBoxI (MYCN^3A^) shows reduced association with RNA ^9^. To evaluate the impact of MYCN-binding to RNA on the proximity of Aurora-A to RNAPII in cells, we performed PLAs between Aurora-A and RNAPII upon HU treatment in cells expressing MYCN^wt^ or MYCN^3A^ (Figure 6A). These assays showed that in the control cells (MYCN^wt^) PLA signals increased slightly between Aurora-A and RNAPII upon replication stress and that this effect was significantly more pronounced in cells expressing MYCN^3A^, arguing that RNA binding of MYCN is required to displace Aurora-A from nascent RNA in cells (Figure 6B, C). To demonstrate that the relocation of MYCN to RNA is essential for removing Aurora-A from RNA, we performed eCLIP experiments in cells expressing MYCN^wt^ or MYCN^3A^. Browser tracks of individual genes showed that binding of Aurora-A to RNA was significantly higher in the MYCN^3A^ condition than in MYCN^wt^ expressing cells (Figure 6D). Transcriptome-wide analyses confirmed this increase in binding, since 9,742 reproducible binding sites of Aurora-A on RNA were identified in SH-EP MYCN^wt^ cells and this number raised to 15,122 enriched windows in SH-EP MYCN^3A^ cells (Figure 6E, F). Comparing the overlap between reproducible Aurora-A windows in MYCN^wt^ and MYCN^3A^ cells revealed that approximately two-thirds of the windows called in MYCN^wt^ were also found in MYCN^3A^ (Figure 6G, H), suggesting that Aurora-A does not relocalize on RNA but rather occupies additional sites. We concluded that Aurora-A accumulates on RNA when MYCN is unable to bind to RNA, arguing that the translocation of MYCN from promoters onto intronic RNA upon stalling of RNAPII displaces Aurora-A from RNA.

**Figure 6:**
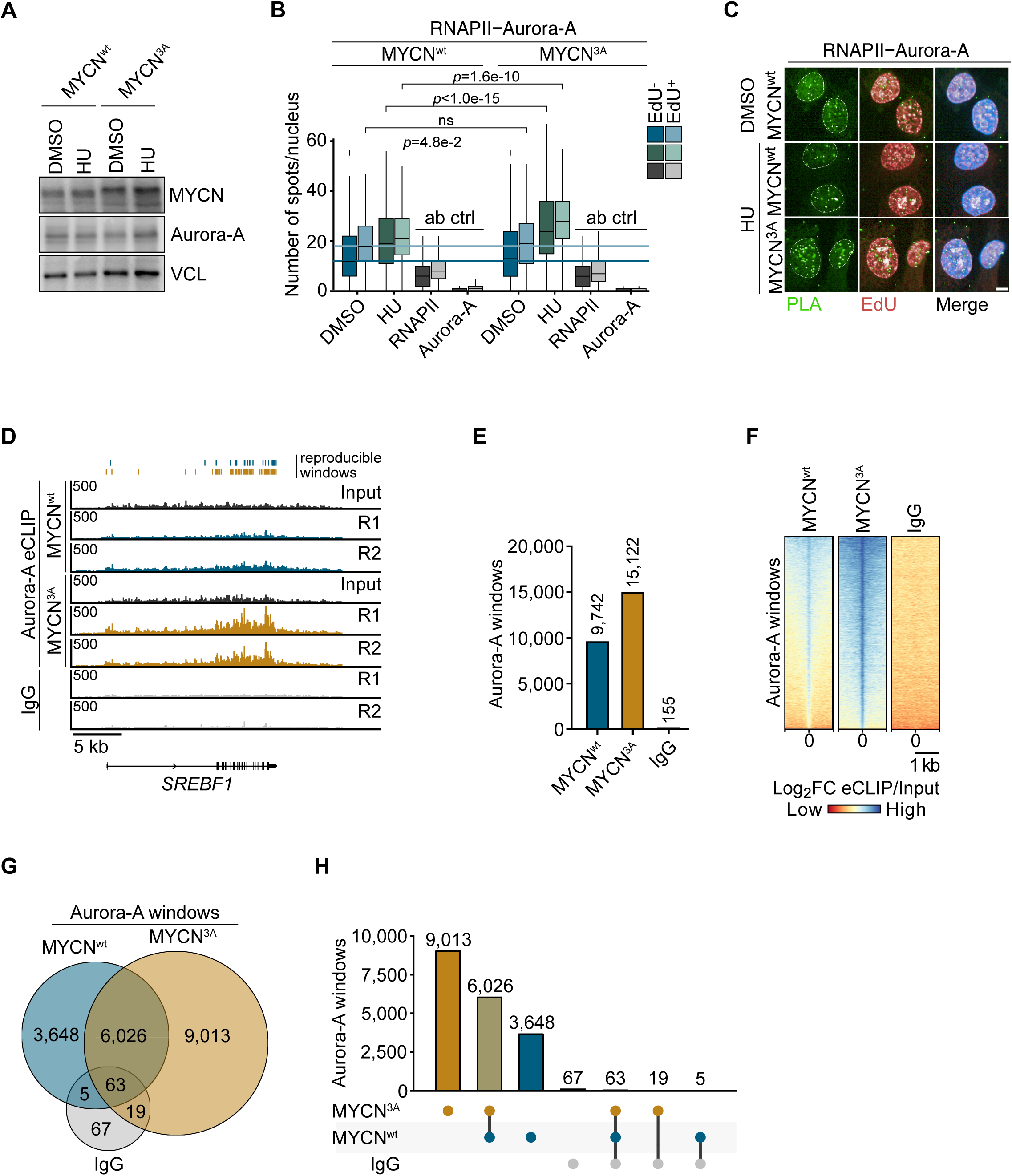
MYCN displaces Aurora-A from RNA. **A.** Immunoblot of indicated proteins in SH-EP MYCN^wt^ or MYCN^3A^ cells after induction of the MYCN construct by doxycycline (1 µg/ml Dox, 24 h) treated with HU (3.5 mM, 2 h) or DMSO as control. VCL was used as a loading control (*n* = 3). **B.** Boxplot of single-cell analysis of nuclear PLA foci between RNAPII and Aurora-A in SH-EP MYCN^wt^ or MYC^3A^ cells treated with HU (3.5 mM, 2 h) or DMSO as control. Conditions are stratified for EdU incorporation. *p* values were calculated comparing the PLA signal of all analyzed cells using an Wilcoxon rank sum test (*n* = 2, ab ctrl = single antibody control). **C.** Representative pictures of PLA foci, EdU staining and the merge with Hoechst described in B. Scale bar indicates 10 µm (*n* = 2). **D.** Read distribution of Aurora-A eCLIPs at the *SREBF1* locus. SH-EP MYCN^wt/3A^ cells were treated for 24 h with doxycycline (1 µg/ml) to induce constructs. Input and IgG are shown as control. Bars on top show reproducible enriched windows (*n* = 3; R = replicate). **E.** Bar graph depicting numbers of reproducible enriched windows of Aurora-A eCLIPs of replicate conditions described in D. Windows in IgG eCLIP are shown as control. **F.** Heatmaps showing the log_2_FC of Aurora-A eCLIPs over the respective size-matched input in depicted conditions sorted for occupancy at merged windows. Shown is one representative replicate per conditions (*n = 3)*. **G.** Venn diagram of reproducible windows of merged replicates of Aurora-A eCLIPs for SH-EP MYCN^wt/3A^ cells. IgG eCLIP as control. **H.** UpSet plot of Aurora-A eCLIP windows comparing enriched windows in SH-EP MYCN^wt^, SH-EP MYCN^3A^ and IgG as a control.

### MYCN-driven tumors are highly dependent on Aurora-A and CDK12

Clinical studies of Aurora-A inhibitors have shown limited efficacy both as monotherapy and in combination with standard chemotherapy ^58–63^. Since the simultaneous inhibition of two proteins that act in a linear pathway can enhance efficacy, we tested whether targeting both Aurora-A and CDK12 increases their effect on MYCN-driven neuroblastoma cells and tumors. Initially, we assessed SR4835 for *in vivo* studies but were unable to achieve sufficient bioavailability; we therefore used the cyclin K degrader PXG-10798, which reduces cyclin K levels at low nanomolar concentrations (DC_50_=0.2 nM, Figure S7A). We tested PXG-10798 side-by-side with SR4835 in neuroblastoma cells and validated that it efficiently degraded cyclin K and moderately reduced CDK12 levels (Figure 7A).

**Figure 7:**
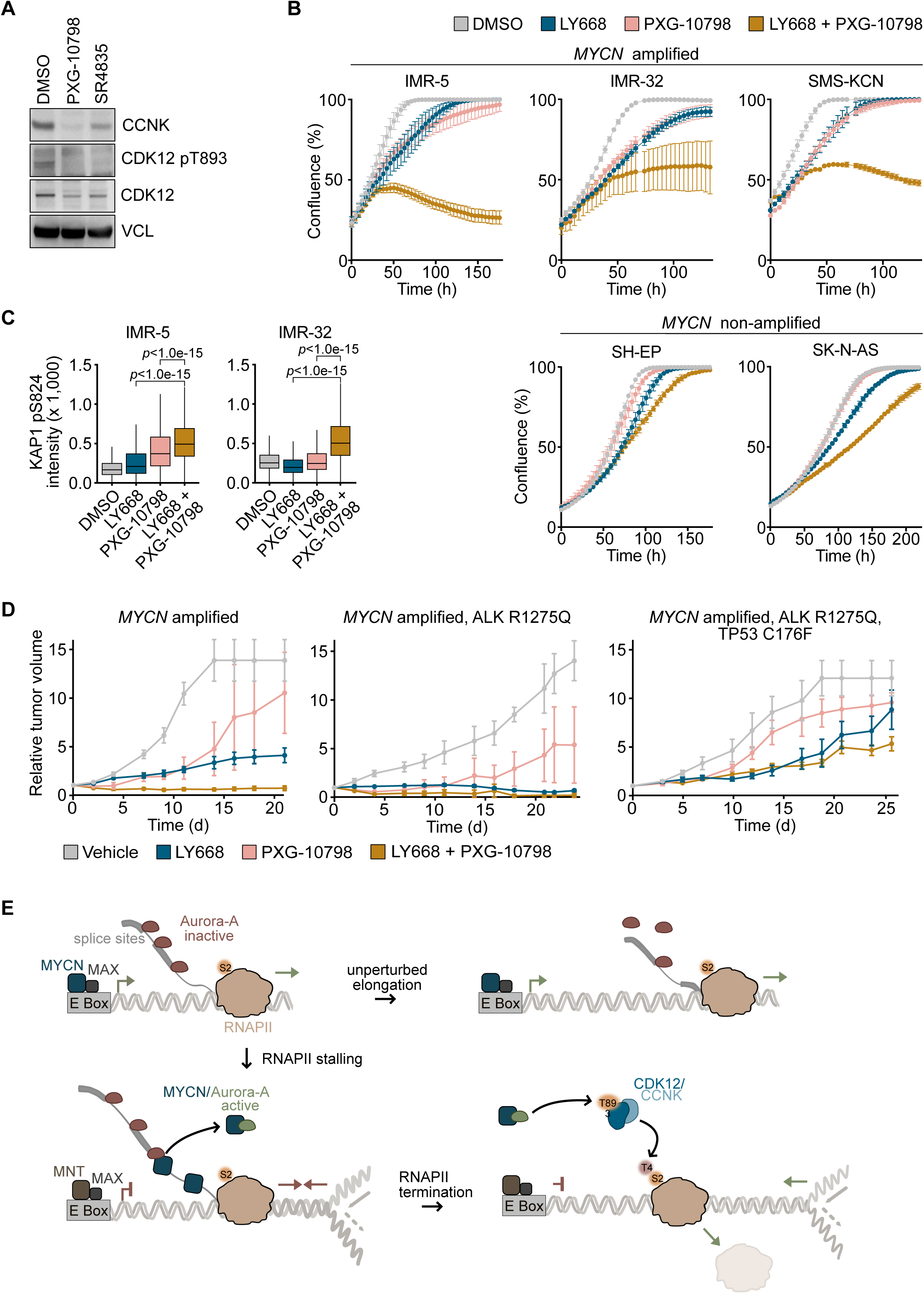
MYCN-driven tumors are dependent on Aurora-A and CDK12. **A.** Immunoblots of indicated proteins in IMR-5 cells upon treatment with SR4835 (100 nM) or PXG-10798 (100 nM) for 6 h. VCL was used as a loading control (*n* = 3). **B.** Growth curves of neuroblastoma cells upon LY668 (30 nM for IMR-5, SH-EP and SK-N-AS; 10 nM for IMR-32; 15 nM for SMS-KCN), PXG-10798 (20 nM for IMR-5, SH-EP and SK-N-AS; 5 nM for IMR-32; 15 nM for SMS-KCN) or the combination as indicated. Data from three technical replicates of one representative experiment is shown as mean ± SD (*n* = 3 for IMR-5, IMR-32, SMS-KCN, SH-EP; *n* = 2 for SK-N-AS). **C.** Boxplot of quantitative immunofluorescence of KAP1 pS824 intensity in IMR-5 (left) and IMR-32 (right) cells treated with LY668 (30 nM for IMR-5; 10 nM for IMR-32), PXG-10798 (20 nM for IMR-5; 10 nM for IMR-32) or the combination for 48 h. *p* values were calculated using Wilcoxon rank sum test. Shown is the single-cell analysis of three technical replicates of one representative experiment (*n* = 3). **D.** Relative changes in tumor volume of three *MYCN* amplified patient-derived xenograft (PDX) models upon treatment as indicated. Tumor volume is normalized to the volume measured at the start of treatment. **E.** Model of our findings. We propose that Aurora-A binds at and close to splice sites on nascent RNA in an inactive form and during unperturbed transcription is removed from nascent RNA upon splicing. Upon stalling of RNAPII, MYCN moves from promoters to nascent RNA, where it complexes with Aurora-A, dislocates Aurora-A from RNA and promotes kinase activation, enabling it to phosphorylate the T-loop of CDK12. Active CDK12 subsequently phosphorylates the CTD of RNAPII at T4, which recruits termination factors to prevent TRCs. See also Figure S7.

Surveying a panel of neuroblastoma cell lines showed that, on average, *MYCN* amplified neuroblastoma cells were more sensitive to PXG-10798 than *MYCN* non-amplified cells (Figures S7B). Combining low concentrations of LY668 with PXG-10798 strongly enhanced growth suppression of three *MYCN* amplified cell lines but had weaker and non-additive effects on two *MYCN* non-amplified cell lines (shown for one combination each in Figure 7B and for four different combinations in IMR-5 and SH-EP cells in Figure S7C). Consistently, immunofluorescence as well as immunoblots showed that phosphorylation of KAP1 was increased when co-treating the cells with low concentrations of LY668 and PXG-10798 (Figure 7C and Figure S7D). To test this regimen in human tumor samples, we treated three *MYCN* amplified patient-derived xenograft models (PDX) of neuroblastoma with LY668, PXG-10798 or the combination of both drugs. Two PDX models also contained an activating ALK mutation and one in addition an inactivating TP53 mutation (Figure 7D). PD studies showed a profound degradation of cyclin K upon single administration *in vivo* (Figure S7E). Combining both inhibitors was more effective than single treatment in all three models tested, causing complete responses in two and a partial response in the TP53-mutant PDX model (Figure 7D). Notably TP53 mutations are very rare in primary neuroblastomas. Collectively, the data argue that the Aurora-A- and CDK12-dependent transcription termination pathway is a specific and targetable dependency of MYCN-driven tumors.

## Discussion

Both MYC and MYCN proteins are transcription factors that affect the expression of a wide range of target genes, suggesting that altering the expression of downstream target genes is the critical oncogenic activity of both proteins. Surprisingly, many effects of MYC and MYCN, such as the ability to suppress the accumulation of promoter-proximal R-loops ^6^ and double-strand breaks ^64^, are observed at all active genes and are independent of changes in gene expression, raising the possibility that gene expression-independent functions of MYC and MYCN contribute to their pervasive oncogenic potential ^65^. Since MYC and MYCN are expressed in a mutually manner in human tumors, the identification of their specific oncogenic functions may open a pathway to selective targeted therapies of *MYCN*-driven tumors.

Here we analyzed a complex of the Aurora-A kinase and MYCN, which forms during the S phase of the cell cycle and is required for S phase progression in *MYCN* amplified neuroblastoma cells ^5,24^. Previous work had implicated Aurora-A in DNA replication ^66^ and in alternate splicing during S phase ^67^, but the critical substrates that enable it to prevent TRCs were previously unknown. Here we established that Aurora-A phosphorylates and activates a pool of CDK12 that is required to prevent TRCs. Activation of CDK12 by Aurora-A is required for phosphorylation of T4, but not S2, in the CTD of RNAPII. While we recapitulate previous data that overall CDK12 activity is required for expression of S phase genes and suppression of intronic polyadenylation ^39,68^, the Aurora-A dependent pool of active CDK12 is not involved in either process but promotes the recruitment of several termination factors. Of these, PCF11 appears to be critical for preventing DNA damage in S phase, but the recruitment of different termination factors may enable RNAPII to respond to the stochastic nature of TRCs, which can occur in either direction and at different locations within a gene ^11,12^.

We found that Aurora-A binds RNA *in vitro* and localizes near splice sites on nascent RNA in cells. RNA binding inhibits Aurora-A kinase, hence RNA-bound Aurora-A has no effect on phosphorylation of CDK12 and RNAPII. We recently showed that RNAPII stalling and the accumulation of intronic RNA causes a fraction of MYCN to accumulate on introns and nascent RNA ^9^. While both recombinant MYCN and Aurora-A bind RNA *in vitro*, the complex of both proteins has a significantly reduced affinity for RNA. Consistently, Aurora-A accumulates on nascent RNA in cells that express a mutant of MYCN that is impaired in RNA binding, suggesting that translocation of MYCN onto RNA removes Aurora-A from nascent RNA in cells and activates its kinase activity to phosphorylate the T-loop of CDK12. Since most nascent, unprocessed RNA is intronic, processes that slow RNAPII elongation will trigger translocation of MYCN onto RNA. We propose that the subsequent displacement of Aurora-A from RNA links the slow-down in elongation to transcription termination and thereby prevents TRCs (see Figure 7E for a model).

These findings have direct implications for the development of Aurora-A-based therapies for MYCN-driven tumors. To date, Aurora-A inhibitors have shown limited therapeutic activity in several tumor types, including neuroblastoma ^58–63^. This is due in part to the fact that clinically available inhibitors have dose-limiting toxicity, which limits their potential therapeutic efficacy. While this may be due in part to inhibition of Aurora-B, the LY668 compound studied here also has shown limited therapeutic activity as a monotherapy and in combination with a backbone chemotherapy regimen containing topotecan, a topoisomerase I inhibitor, and cyclophosphamide, a DNA alkylating agent ^63^. Our data show that MYCN/Aurora-A complexes coordinate transcription with DNA replication via activation of CDK12. Inhibition of Aurora-A with a cyclin K degrader that inhibits CDK12 shows strong additive effects in causing DNA damage and suppressing MYCN-driven neuroblastoma growth. Since both molecules disrupt the coordination of transcription with DNA replication during the S phase of the cell cycle, the ensuing DNA damage depends on ongoing DNA and RNA synthesis. In contrast, the topoisomerase inhibitors and DNA alkylating agents that are used in chemotherapy arrest the cell cycle via induction of p53, often in the G1 phase of the cell cycle, and inhibit both transcription and replication, so these compounds directly counteract the effects of Aurora-A and CDK12 inhibition. The data presented here therefore provide a rationale for exploring combination therapies of Aurora-A and CDK12 inhibitors in MYCN-driven tumors without the confounding effects of conventional chemotherapeutic agents.

## Acknowledgements

This work was supported by grants from the Mildred Scheel Junior Research Center Program (#70113303 to G.B.), the European Research Council (SENATR ERC #101096948 to M.E.), the German Cancer Aid (#70114538 to M.E and #70114008 to M.G.), the German Research Foundation (CRC1588 #493872418 to G.B., INST 93/1023-1-FUGG to M.E.), Cancer Research UK (C24461/A23303 to R.B.), the Medical Research Council (MR/X008673/1 to R.B.), UKRI (BB/X002780/1 to P.A.E., D.P.B and C.E.E), and the Alex’s Lemonade Stand Foundation Crazy 8 Initiative (to G.B. and M.E.).

We thank Carsten Ade for performing the sequencing experiments, Peter Gallant for help with bioinformatic analyses, Tobias Roth and André Kutschke for technical help and Dennis Gürgen for help with animal experiments.

## Author Contributions

Conceptualization, G.B. and M.E.; Methodology, G.B. and M.E.; Investigation, M.M., L.E., D.F., N.G., S.F., M.S., I.H.K., S.G.B., I.K., D.P.B., S.A.H., C.S.-V., V.N.-P. and G.B.; Formal Analysis, D.S., R.V., I.K. and L.U.; Visualization, M.M. and G.B.; Writing – Original Draft, G.B. and M.E.; Supervision, P.A.E., C.E.E., M.G., D.P., M.B., M.W.R, R.B., M.E. and G.B.; Funding Acquisition, G.B., R.B., M.G., P.A.E., D.P.B., C.E.E. and M.E.

## Supplemental information

Document S1: Figures S1-S7 and Table S1

## STAR Methods

### CONTACT FOR REAGENT AND RESOURCE SHARING

Further information and requests for resources and reagents should be directed to and will be fulfilled by the Lead Contact, Gabriele Büchel.

### EXPERIMENTAL MODEL AND SUBJECT DETAILS

#### Cell culture

Neuroblastoma cell lines (IMR-5, NHO2A, IMR-32, SMS-KCN, SK-N-BE, SH-EP, SH-SY5Y and S-KN-AS) and primary cell lines were grown in RPMI 1640 medium (Thermo Fisher Scientific). HEK293TN cells were grown in DMEM (Thermo Fisher Scientific). Medium was supplemented with 10% fetal calf serum (Biochrom) and penicillin-streptomycin (Sigma-Aldrich). SH-EP MYCN^wt^ and MYCN^3A^ cells were kindly provided from ^9^.

All cells were routinely tested for mycoplasma contamination. Where indicated, cells were treated with LY-3295668 (1 µM, Selleckchem/Biozol), SR-4835 (50 nM, Axon Medchem), PXG-10798 (100 nM, Proxygen) Hydroxyurea (3.5 mM Sigma-Aldrich), Irinotecan (20 µM, Sigma-Aldrich), CBL0137 (5 µM, Selleckchem/Biozol), Cisplatin (5 µM, Medchem), Aphidicolin (1 µM, Abcam), Thymidine (2 mM, Sigma-Aldrich) or DMSO (Sigma-Aldrich).

For live growth imaging, cells were seeded in 24-96 well plates (Greiner) and treated as indicated. Cell growth was analyzed up to 7 days, monitored by phase-contrast confluence by time lapse microscopy using the Incucyte SX5 system, with at least 4 fields measured every 6 h using 10x magnification.

### METHOD DETAILS

#### Cloning

To express Aurora-A^wt^, Aurora-A^T208D^ or Aurora-A^T208E^ double-stranded DNA fragments (gBlock™ User Guide, IDT) based on cDNA sequences of Aurora-A with or without respective point mutation were cloned into the pRRL vector containing an SFFV promoter. For transformation of the engineered plasmids, XL-1 blue *E. coli* cells were used. The integrity of the DNA was confirmed by Sanger sequencing (LGC genomics).

#### Transfection and lentiviral infection

For lentivirus production, HEK293TN cells were transfected using PEI (Sigma-Aldrich). Engineered plasmids were transfected together with the packaging plasmid psPAX.2 and the envelope plasmid pMD2.G. Virus-containing supernatant was collected 48 h and 72 h after transfection. NHO2A cells were infected with lentiviral supernatant in the presence of 4 µg/ml polybrene for 24-48 h. Infected cells were selected with puromycin.

IMR-5 cells were transfected with siRNAs using Lipofectamine RNAiMAX (Thermo Fisher Scientific) on a 96-well plate. Experiment was analyzed 72 h post siRNA transfection.

#### Immunoblot and Immunoprecipitation

Cells were harvested by centrifugation followed by lysis in RIPA buffer (50 mM HEPES pH 7.9, 140 mM NaCl, 1 mM EDTA, 1% Triton X-100, 0.1% SDS, 0.1% sodium deoxycholate) containing protease and phosphatase inhibitors (Sigma-Aldrich) and incubated for 30 min on ice. The lysate was cleared by centrifugation and the protein concentration was determined by Bradford assay. Cell lysate was separated by Bis-Tris-PAGE and transferred to PVDF membranes (Millipore). Membranes were blocked for 1 h with BSA in TBS-T and incubated using the indicated antibodies overnight at 4°C. Antibodies used are listed in Key Resources Table. As loading control, vinculin (VCL) or actin (ACTB) were used. The day after, membranes were washed and probed for 1 h at RT with HRP-conjugated secondary antibodies. Images were acquired using the Fusion FX7 EDGE imaging system (Vilber).

For co-immunoprecipitations, depending on the antibody species, 20 µl per IP of A, G or a 1:1 A/G Dynabeads mix (Thermo Fisher Scientific) was washed with 5 mg/ml BSA-PBS and incubated overnight at 4°C with 2.5 µg antibody against the protein of interest. Harvested cells were lysed with HEPES lysis buffer (20 mM HEPES pH 7.9, 150 mM NaCl, 0.2% v/v NP-40, 0.5 mM EDTA, 10% v/v Glycerol, 2 mM MgCl_2_), briefly sonicated, incubated with 50 U Benzonase (Merck) for 1 h at 4°C, then 2-4 mg of lysate was added to the bead/antibody mix and incubated for 6-8 h at 4°C. 1-2% of the lysate was kept as input reference. Lysates were eluted in 2x Laemmli, after washing with HEPES lysis buffer, boiled for 5 min at 95°C and subsequently subjected to immunoblotting.

#### Sucrose gradient fractionation and precipitation

Whole cell lysates were loaded on a 10-50% sucrose gradient and separated by ultracentrifugation in a Beckman Optima L-90 LK ultracentrifuge (Beckman Coulter) for 18 h at 30,000 rpm and 4°C using a SW41 rotor (Beckman Coulter). The fractions were obtained by manual fractionation by removing 500 µl off the top of the gradients, then the fractions were subjected to co-immunoprecipitation.

#### Protein digestion and TMT labelling

10*10^6^ NHO2A Aurora-A^wt^ or Aurora-A^T208D^ cells were harvested after 8 h treatment with 1 µM LY668 or DMSO as control. Cells were resuspended in lysis buffer (50 mM HEPES pH 8, 1% SDS, 1% Triton X-100, 1% NP-40 (IGEPAL), 1% Tween-20, 1% SDC, 50 mM NaCl, 1% glycerol, 1x cOmplete EDTA-free protease inhibitors (Roche), 1x PhosStop phosphatase inhibitors (Roche)) were sonicated three times for 30 s, and then heated to 80°C for 5 min. Following centrifugation at 17,200 x*g*, solubilised protein extracts were treated with 4 mM DTT at 60°C for 10 min, allowed to cool to RT and alkylated with 14 mM IAA (30 min, RT, in the dark), and quenched by addition of DTT (7 mM final). Proteins were then precipitated onto 1:1 hydrophilic and hydrophobic Sera-Mag magnetic beads (GE healthcare) in 70% (v/v) ethanol, followed by three washes in 80% (v/v) ethanol. On-bead proteolytic digestion was performed in 100 mM TEAB using Trypsin Gold (Promega) at 1:50 (w/w) ratio, at 37°C overnight. Peptide concentration was estimated using Pierce^TM^ peptide colorimetric assay (ThermoFisher Scientific) following the manufacturer’s instructions, and 100 µg of protein from each sample was labelled with TMT (ThermoFisher Scientific) as per manufacturer’s instructions. Three TMT-10 multiplexing sets were used, where each set was assigned to one biological replicate from each of the 9 conditions, and a pooled sample was prepared for use as a common reference between the sets during data analysis. Peptides were incubated with TMT reagents for 1 h, and labelling was quenched with 5% hydroxylamine at 0.3% final concentration for 15 min. Labelled peptides from each TMT set were then pooled together and dried to completion by vacuum centrifugation.

#### Reverse-phase fractionation

Each set of TMT-labelled peptides was fractionated by high-pH reversed-phase (RP) HPLC over a 60 min gradient as previously described ^70^. In brief, multiplexed labelled samples were resuspended in 94.5% (v/v) solution A (20 mM NH_4_OH, pH 10) and 5.5% (v/v) solution B (20 mM NH_4_OH in 90% v/v acetonitrile). Fractionation was performed on an Extend C18 (3.5 μm, 3 mm x 150 mm) Agilent column. Peptides were eluted with an increasing concentration of solution B up to 30% over 25 min, then 75% B over 12 min, at a flow rate of 0.5 ml/min for 1 h. Sixty-one fractions were collected and concatenated to 11 pools. For each pool, 5% was aliquoted and dried to completion prior to MS analysis. The remaining 95% was subjected to TiO_2_-based phosphopeptide enrichment, as described previously ^71,72^.

#### Phosphopeptide enrichment

TMT-labelled peptides were resuspended in loading buffer (80% ACN, 5% TFA, 1 M glycolic acid) and mixed with TiO_2_ beads (Titansphere, GL Sciences) for 20 min with shaking, followed by three sequential washes, first with loading buffer, then with 80% ACN/1% TFA, and finally with 10% ACN/0.2% TFA. Phosphopeptides were eluted in two consecutive incubation steps with 1% and 5% ammonium hydroxide, for 10 min each. Peptides were dried to completion and resuspended in 3% ACN/0.1% TFA prior to LC-MS/MS analysis.

#### LC-MS/MS analysis

Global proteomics and phosphopeptidomics of TMT-labelled peptides was carried out by liquid chromatography-tandem mass spectrometry (LC-MS/MS) using a nanoflow Ultimate3000 LC system (Thermo Scientific) coupled online to an Orbitrap Fusion Lumos (Thermo Scientific). Peptides were first loaded onto a PepMap100 trapping column (Thermo Scientific) packed with C18 resin (300 µm x 5mm), for 7 min with a flow rate of 12 µl/min. Peptides were then separated on a 50 cm x 75 µm EasySpray column (Thermo Scientific) with 2 µm C18 particles over a 90 min linear gradient from 3.8% to 50% solution B (80% ACN/0.1% FA), where solution A was 0.1% FA. Column temperature was set to 45 °C. The mass spectrometer was interfaced with a FAIMS Pro Duo device (Thermo Scientific) with a static carrier gas flow of 3.7 l/min, operating with a standard resolution and cycling between three compensation voltages (−40, - 55, and -70) for the full duration of the acquisition method. Data was acquired in data-dependent mode with a constant 1.5 s cycle time. Precursors were sampled over *m/z* range of 400-1600, with an Orbitrap resolution of 120,000, a standard AGC target, and an automatically determined IT. Fragments with *m/z* above 120 were isolated from a 0.7 *m/z* window and resolved in the Orbitrap at 50,000, using 38% HCD normalised collision energy, 200% AGC target, and 200 ms max IT.

#### In vitro protein expression and purification

For IC_50_ determinations, full length mouse (Uniprot Q7TNK2) Aurora-A, or constructs encoding Asp or Glu point mutations at position T208 (mouse) cDNAs, were sub-cloned into N-terminal 6His-tag encoding pET28a and after IPTG-induced expression in BL21(DE3) pLysS strain, purified to homogeneity using Ni-affinity chromatography and size exclusion chromatography. The recombinant Aurora-A kinase domain (amino acids 122-403, 35.8 kDa) with three-point mutations C290A, K339C, C393A was expressed in *Escherichia coli* BL21 (DE3) pLysS strain using a pETM11 bacterial expression vector with a His-TEV construct. The synthetic MYCN peptide (amino acids 28-89, 7.3 kDa) with a reactive maleimide group was ordered from PeptideSynthetics (UK). Chemical crosslinking was performed with 2-fold molar excess of MYCN peptide over reduced Aurora-A kinase domain in crosslinking buffer (200 mM NaCl, 100 mM K_2_HPO_4_ (pH 7), 10% glycerol, 5 mM EDTA) overnight at room-temperature. Proteins were purified by immobilized metal affinity chromatography followed by size-exclusion chromatography. All proteins were stored in storage buffer (200 mM NaCl, 100 mM K_2_HPO_4_ (pH 7), 10% glycerol, 5 mM MgCl_2_).

For CDK12/cyclin K e*xpression* the CDK12 kinase domain (amino acids 714-1063) and the cyclin K cyclin box domains (amino acids 1-267) were synthesized as expression-optimized genes by GeneArt. These genes were cloned into a pACEBac1 acceptor vector, modified to include an N-terminal GST-affinity tag followed by a tobacco etch virus (TEV) protease cleavage site, respectively. Vectors were used for co-expression in *Sf9* insect cells utilizing the MultiBac^Turbo^ system ^73^.^73^. The CDK12/CycK complex was purified using a combination of GST-affinity chromatography with sequential size-exclusion chromatography as described in ^74^. Expression constructs for the Cdk12 T893A point mutation were generated using site-directed mutagenesis. The mutant Cdk12 was then co-expressed with cyclin K in Sf9 insect cells and purified following the same protocol used for the wild-type protein.

For co-expression experiments, wild-type or mutant CDK12 kinase domain constructs as well as cyclin box constructs of cyclin K were inserted into pACEBac1 acceptor vectors as described above. Additionally, either full-length CDK-activating kinase CAK1 from *S. cerevisiae* or Aurora-A kinase domain (amino acids 122-403) with three-point mutations C290A, K339C, C393A was also inserted into an unmodified pACEBac1 acceptor vector without affinity tags. CDK12/cyclin K vectors were then co-expressed in *Sf9* insect cells using the MultiBac^Turbo^ system in absence or presence of either CAK1 or Aurora A constructs. Purification of CDK12/cyclin K constructs co-expressed with CDK-activating kinases was conducted as described above for the wild-type constructs0.2. Purified CDK12/cyclin K complexes were subjected to radioactive kinase activity assays or western blot experiments using a phospho- and site-specific CDK12 [α-pT893] antibody.

For RNAPII GST-CTD_[52]_ expression the wild-type C-terminal domain of RNAPII (residues 1587–1970) was cloned into a pGEX-6P1 vector modified to include an N-terminal GST tag and a 3C protease cleavage site. Expression of GST-CTD52 was induced with IPTG and carried out overnight at 20°C. Purification involved GST-affinity chromatography followed by size-exclusion chromatography on a Superdex 200 column, with the GST tag retained.

The expression constructs, pET30TEV Aurora-A 1-403 ^21^, pETM6T1 3xFLAG-MYCN 1-137 ^21^ and pCDF lambda phosphatase ^75^ were generated in previous work. Unphosphorylated Aurora-A 1-403 protein was produced by co-expression with lambda phosphatase. Kinase expression and purification was undertaken as in ^56^. Expression and purification of 3xFLAG-MYCN 1-137 was carried out as described in earlier work ^21^.

#### Electrophoretic mobility shift assay

Recombinant Aurora-A kinase domain or crosslink (Aurora-A:MYCN) was incubated with 5 nM Cy5-labeled UG-High RNA purchased from GenScript (Netherlands, sequence Cy5-UGC UGG GAU GGU CAU GGU GGG UGA CCC UGG GAU GGC CGC GGU GGG UGA CCC UGG AAU GGU) for 30 min at room-temperature in protein storage buffer. 2 μl of 80% glycerol was added prior to loading 10 μl of the binding mixture onto NativePAGE. The complexes were separated at 100 V at 4°C for 90 min. Cy5 fluorescence was detected using Licor Imaging Systems (Odyssey). Each EMSA was performed in five independent replicates. The percent of free-RNA probe was quantified using ImageJ and plotted using Prism version (v8.2.1).

#### In vitro kinase assays

For IC_50_ determinations, full length murine Aurora-A cDNA, or the indicated point mutations, were cloned into pET28a and purified and substrate phosphorylation was measured in real-time using a linearized non-radioactive substrate mobility shift assay, as previously described ^76^.

For Aurora-A/CDK12 assays, 0.5 µM of recombinant Aurora-A (122-403, C290A, K339C, C393A), 1 µM MYCN (1-123) and 1 µM CDK12 (714-1063)/CyclinK (1-267) were treated with either 5 µM UG-rich RNA or 0.5 µM of the Aurora-A inhibitor LY-3295668. Reactions were carried out in kinase buffer (20 mM Tris pH 7.5, 25 mM NaCl, 1 mM MgCl_2_, 1 mM DTT, 0.01% TWEEN 20). The reaction was started by addition of ^32^P-ATP (Hartmann Analytics) and was stopped by addition of 1x volume of 2x Laemmli buffer. Samples were analyzed by SDS-PAGE followed by autoradiography. Statistical analysis was carried out in Prism (Graphpad Software, Inc.).

#### [^32^P]-γ-ATP kinase activity assay

Radioactive kinase activity assays were performed using 0.5 µM indicated recombinant kinase in kinase buffer (50 mM HEPES pH 7.6, 34 mM KCl, 7 mM MgCl_2_, 2.5 mM DTE, 5 mM β-glycerophosphate) supplemented with 0.2 mM ATP containing 0.45 mCi [^32^P]-γ-ATP/ml (PerkinElmer) in a combined volume of 15 µl per sample. To initiate the kinase reaction, 100 µM pS7-CTD_[3]_ peptide (three CTD repeats with serine at position 7 pre-phosphorylated, purchased from Biosynthan, Berlin) was added. The kinase reaction was incubated at 30°C and 300 rpm for 1 h. To terminate the kinase reaction, EDTA to a final concentration of 50 mM was added. Subsequently, the reaction samples were individually spotted onto Amersham Protran nitrocellulose membranes (Cytiva) and subjected to three washing steps of 5 min each with 0.75% (v/v) phosphoric acid. These membranes were then transferred to 6 ml liquid scintillation vials (Sarstedt), and 2 ml liquid scintillation cocktail Ultima Gold (PerkinElmer) was added to allow for radioactive count determination on a liquid scintillation counter LS 6500 (Beckman Coulter) for a duration of 1 min per sample. Measurements were performed in triplicate and depicted as mean with standard deviation (SD).

#### ATP kinase activity assay

Kinase assays were conducted using 0.2 µM CDK12/cyclin K recombinant kinase complex or Aurora A kinase in kinase buffer (50 mM HEPES pH 7.6, 34 mM KCl, 7 mM MgCl_2_, 2.5 mM DTE, 5 mM β-glycerophosphate) containing 0.2 mM ATP. Kinase reactions were initiated by the addition of 10 µM GST-CTD_[52]_, incubated at 30°C and 300 rpm, and terminated at indicated time points by the addition of equal volumes of 4x SDS sample buffer. If indicated, SR-4835 was pre-incubated with the respective kinase at a final concentration of 1 mM for 5 min prior to substrate addition. Samples were analyzed by SDS-PAGE or western blotting using indicated CTD phosphorylation specific antibodies.

#### ADP-Glo kinase assays

For assays using unphosphorylated Aurora-A 1-403 and 3xFLAG-MYCN 1-137, reaction mixes were made containing 0.5 μM unphosphorylated Aurora-A 1-403, 40 μM myelin basic protein (Sigma-Aldrich), and a 2-fold serial dilution of 2.5 μM 3xFLAG-MYCN 1-137 in a final reaction buffer of 20 mM Tris pH 7.5, 25 mM NaCl, 10 mM MgCl_2_, 1 mM DTT, 0.01% TWEEN 20, 1 U/μl murine RNase inhibitor (NEB). Kinase activity was initiated by the addition of 40 μM ultra pure ATP (Promega) in a final reaction volume of 10 μl. Reactions were incubated at RT for 45 min in white, opaque 96-well ½ area plates (PerkinElmer) with gentle shaking. 10 μl ADP-Glo reagent (Promega) was used to terminate the assay and the plate incubated at RT for 40 min with gentle shaking to remove any residual ATP. 20 μl ADP-Glo detection reagent (Promega) was then added, and the plate incubated for a further 40 min at RT with gentle shaking. Reactions were performed in triplicate and luminescence was read on a Victor X5 plate reader (PerkinElmer). Background signal, measured from reactions set up without ATP, were subtracted from the data for each experiment. Data were analyzed in Graphpad Prism (version 10.3.1) and fitted by nonlinear regression to a four-parameter dose-response stimulation function.

#### In vitro co-precipitation assay

Biotinylated UG^high^ RNA oligonucleotide, 5’-bio-UGC UGG GAU GGU CAU GGU GGG UGA CCC UGG GAU GGC CGC GGU GGG UGA CCC UGG AAU GGU-3′ was synthesized by IDT.

5 μM biotinylated UG^high^ RNA oligonucleotide was incubated with 6 μM selected interactor(s) for 2 h at 4^°^C in the assay buffer, 50 mM Tris pH 7.5, 150 mM NaCl, 5 mM MgCl_2_, 5 mM β-mercaptoethanol, 0.02% TWEEN-20, 1 U/μl murine RNase inhibitor (NEB). The reaction mixes were then added to 20 μl Streptavidin Sepharose beads (IBA) equilibrated in assay buffer for 30 min at 4^°^C and washed three times with 900 μl assay buffer. 40 μl SDS-loading buffer was added to each reaction and boiled prior to SDS-PAGE analysis.

For co-precipitation assays in which the concentration of 3xFLAG-MYCN 1-137 was varied, reactions were carried out as described above, but 3xFLAG-MYCN 1-137 protein was included at concentrations 0-30 μM, to create assay mixes with molar ratios of unphosphorylated Aurora-A 1-403: 3xFLAG-MYCN 1-137 of 1:0, 1:0.5, 1:1, 1:2 and 1:5.

#### Spike-in chromatin immunoprecipitation sequencing

ChIP and ChIP-Rx were performed as described previously ^25^. In brief, for each ChIP-Rx experiment, 5 to 8*10^7^ cells per immunoprecipitation were harvested, proteins were crosslinked to chromatin for 5 min at RT with formaldehyde (final concentration 1%). The reaction was quenched by addition of 125 mM Glycine for 5 min at RT. Cell pellets were resuspended in PIPES lysis buffer containing protease and phosphatase inhibitors (5 mM PIPES pH 8.0, 85 mM KCl, 0.5% v/v NP-40) and IMR-5 ChIP-Rx experiments were spiked with murine NHO2A cells (ratio 1:10). Nuclei were incubated for 20 min on ice, pelleted by centrifuging, then resuspended in RIPA lysis buffer (50 mM HEPES pH 7.9, 140 mM NaCl, 1 mM EDTA, 0.1% v/v SDS, 1% v/v Triton X-100, 0.1% w/v sodium deoxycholate) and incubated for 10 min at 4°C. Chromatin was fragmented using the Covaris M220 Focused Ultrasonicator (Peak Power = 75.0, Cycles/Burst = 200, Duty Factor = 10.0, Duration = 3,000 s per ml cell lysate for IMR-5 cells, Duration = 5,400 s per ml cell lysate for NHO2A cells, < 3*10^7^ cells/ml). Sonicated chromatin size of 150-200 bp was verified by agarose gel electrophoresis. 1% of the lysate was kept for input reference. A total volume of 30 µl (for ChIP) or 100 µl (for ChIP-Rx) per IP of A, G or 1:1 A/G Dynabeads mix (Thermo Fisher Scientific) was washed with BSA-PBS and incubated with 3-5 µg (for ChIP) or 15-20 µg (for ChIP-Rx) antibody against RNAPII (A-10), pS2 RNAPII, pT4 RNAPII or pH3S10 overnight at 4°C on a rotating wheel. Chromatin samples were cleared by centrifugation, then incubated with the antibody/beads mix for 6 h at 4°C on a rotating wheel. Samples were washed three times each with washing buffer I (20 mM Tris-HCl pH 8.1, 150 mM NaCl, 2 mM EDTA, 1% v/v Triton X-100, 0.1% v/v SDS), washing buffer II (20 mM Tris-HCl pH 8.1, 500 mM NaCl, 2 mM EDTA, 1% v/v Triton X-100, 0.1% v/v SDS), washing buffer III (10 mM Tris pH 8.1, 250 mM LiCl, 1 mM EDTA, 1% v/v NP-40, 1% w/v sodium deoxycholate) and twice with TE buffer. The elution of the chromatin was performed by incubating the beads with elution buffer (100 mM NaHCO_3_, 1% v/v SDS in TE buffer) for 15 min at RT on a rotating wheel and repeated for a total of two elution steps. Input and immunoprecipitation samples were de-crosslinked by digestion with RNase A for 1 h at 37°C and followed by incubation overnight at 65°C with shaking, and digestion with proteinase K for 2 h at 45°C. The isolation of the DNA was performed by phenol-chloroform extraction, purified with ethanol precipitation and subsequently quantified with the Quant-iT PicoGreen dsDNA assay (Thermo Fisher Scientific) for ChIP-Rx experiments. Manual ChIP samples were subjected to RT-qPCR (see Key Resources Table for primer sequences). DNA libraries of ChIP-Rx samples were prepared by using the NEBNext Ultra II DNA Library Prep Kit for Illumina Sequencing (New England Biolabs).

#### Total RNA sequencing

RNA sequencing was performed by extracting RNA with the RNeasy Mini Kit (Qiagen) according to the manufacturer’s instructions. Total RNA was purified using the NEBNext rRNA depletion kit (NEB). RNA concentration and RQN values were determined with a Fragment Analyzer (Agilent) and 1 ng RNA with RQN > 9 was used for library preparation using the Ultra II Directional RNA Library Prep for Illumina following the manufacturer’s manual. Libraries were size selected using SPRIselect Beads (Beckman Coulter) after amplification with 12 PCR cycles. Library quantification and size determination were performed with the Fragment Analyzer (Agilent) using the NGS Fragment High Sensitivity Analysis Kit (1-6,000 bp; Agilent). Libraries were subjected to cluster generation and base calling for 100 cycles paired end on the Illumina NextSeq 2000 platform.

#### (Single-End) enhanced crosslinking and immunoprecipitation

eCLIP and seCLIP were performed as described in ^9,53,77^. All eCLIP experiments were performed in biological duplicates or triplicates with one corresponding pooled size-matched input using 40*10^6^ cells per replicate and condition. Cells were crosslinked with UV light at 254 nm for 2 min at 400 mJ/cm^2^. IgG eCLIPs were performed as controls. Cells were lysed in cold lysis buffer (50 mM Tris-HCl pH 7.4, 100 mM NaCl, 1% v/v Igepal CA-630, 0.1% v/v SDS and 0.5% w/v sodium deoxycholate) supplemented with protease inhibitors and murine RNase inhibitor (New England Biolabs). Afterwards lysates were sonicated with a M220 Focused-ultrasonicator (Covaris) for 3 min at low settings and treated with Turbo DNase and RNase I. Lysates were cleared with centrifugation, then the supernatant was submitted to A/G Dynabeads (Thermo Fischer Scientific) pre-coupled with 10 µg Aurora-A antibody (Abcam) or murine IgG Control Antibodies (Sigma-Aldrich). After overnight incubation, 2% v/v lysate was kept as input and samples washed with wash buffer (20 mM Tris-HCl pH 7.4, 10 mM MgCl_2_, 0.2% v/v Tween-20 and 5 mM NaCl) and high-salt wash buffer (50 mM Tris-HCl pH 7.4, 1 M NaCl, 1% v/v Igepal CA-630, 0.1% v/v SDS and 0.5% w/v sodium deoxycholate). RNA was dephosphorylated, then 3‘ RNA adapter ligation was performed using T4 RNA ligase (New England Biolabs) and A01, B06 and D08 barcoded RNA adaptors ^53^ for eCLIPs (samples were multiplexed afterwards) and InvRiL19 RNA adapter for seCLIPs ^77^ (sequences in Key Resources Table). Samples were separated on 10% Bis-Tris gels and transferred to nitrocellulose membranes (Merck) overnight at 4°C at 30 V. Samples were cut at the observed protein size extending above, then incubated with proteinase K (New England Biolabs), followed by RNA purification using the RNA Clean and Concentrator-5 kit (Zymo Research). Input samples were dephosphorylated and 3’ adaptors were ligated before samples were reverse transcribed using the AR17 primer for eCLIP and InvAR17 for seCLIP and the AffinityScript enzyme (Agilent). Samples were purified using MyOne Silane Dynabeads (Thermo Fisher Scientific), then 5’ adapter ligation was performed with Rand3Tr3 for eCLIP and Invrand3Tr3 for seCLIP. Optimal number of PCR cycles was determined via qPCR using the PowerUP SYBR Green Master Mix (Thermo Fisher Scientific) and Illumina compatible primers. Cycle estimation was run at a StepOnePlus Real-Time PCR System (Thermo Fisher Scientific) and optimal PCR cycles were determined from C_t_ values. All samples were amplified with the Q5 High-Fidelity 2x PCR Master Mix (New England Biolabs) and size selected with SPRIselect beads (Beckman Coulter). The Aurora-A seCLIP in IMR-5 cells was spiked with cDNA for normalization before PCR amplification. The Aurora-A eCLIP in SH-EP MYCN^wt/3A^ cells was multiplexed after 3‘ RNA adapter ligation using individual barcodes.

#### Immunofluorescence

Cells were seeded into 96-well plates (Perkin Elmer/Revvity) and treated as indicated with inhibitors or siRNA transfection, 1 h before fixation cells were pulsed with 10 µM EdU (Jena Bioscience). Cells were fixed with 4% PFA for 10 min and permeabilized with 0.3% Triton X-100 for 10 min at RT or fixed/permeabilized with 100% Methanol on ice for 20 min and blocked with 5% BSA-PBS for 30 min at RT. Newly synthesized DNA was visualized using click chemistry by performing a copper(I)-catalyzed azide-alkyne cycloaddition (100 mM Tris pH 8.5, 4 mM CuSO4, 10 mM AFDye 647 Azide (Jena Bioscience), 10 mM L-Ascorbic Acid). Cells were incubated with primary antibodies in blocking buffer overnight at 4°C, then incubated with fluorophore-conjugated secondary antibodies for 1 h at RT with prior and post washing steps with PBS. Nuclei were counterstained with Hoechst 33342 (2.5 µg/ml, Sigma) for 10 min at RT.

Image acquisition was performed by using the Operetta CLS High-Content Analysis System (Revvity) with 40-63x magnification and images were processed using the Harmony High Content Imaging and Analysis Software (Revvity) and R. Cells were grouped into cell cycle phases according to the EdU and Hoechst content of the control conditions.

#### In situ Proximity Ligation Assay

3,500-7,000 IMR-5 cells were seeded per well in a 384 well format (PerkinElmer). Cells were treated with inhibitors as indicated and if cells were differentiated according to cell cycle phases, conditions were pulsed with 10 µM EdU (Jena Bioscience). For SH-EP MYCN^wt^/MYCN^3A^ 750 cells were seeded in a 384 well format and constructs were induced with 1 µg/ml doxycycline (Sigma-Aldrich) for 24 h. Cells were fixed/permeabilized with 100% methanol (Merck) on ice for 20 min. PLA was performed according to manufacturer’s instructions (Duolink *In Situ* Proximity Ligation Assay, Merck). After blocking with blocking solution for 30 min cells were incubated with primary antibodies in blocking solution overnight at 4°C. After washing with PBS, PLUS (anti-rabbit) and MINUS (anti-mouse) probes were conjugated to primary antibodies for 1 h at 37°C. The cells were washed with Wash Buffer ‘A’ and the ligation reaction was carried out for 30 min at 37°C, followed by washing and *in situ* PCR amplification for 2 h at 37°C. Before washing with Wash Buffer ‘B’ EdU was visualized using click chemistry as described for immunofluorescence and nuclei were counterstained with Hoechst33342 (Thermo Fisher Scientific). Image acquisition was performed using the Operetta CLS High-Content Analysis System with 40x magnification (Revvity) and images were processed using the Harmony High Content Imaging and Analysis Software (Revvity). Single cell results were analyzed, and *p*-values were calculated with RStudio.

#### Flow cytometry

For flow cytometry analysis subconfluent cells were labelled with 20 µM 5-Bromo-2’-deoxyuridine (BrdU, Sigma-Aldrich) for 1 h. Cells were harvested, washed with ice-cold PBS and fixed in 80% ethanol overnight at -20°C. Cells were washed with cold PBS and incubated in 2 M HCl/0.5% Triton X-100 for 30 min at room temperature. Cell pellets were neutralized by incubating with Na_2_B_4_O_7_. The pellet was incubated with Anti-BrdU-FITC antibody (BioLegend) diluted in 100 μl 1% BSA, 0.5% Tween-20 in PBS for 30 minutes at room temperature in the dark. After washing with PBS, the cells were re-suspended in PBS with RNase A (24 µg/ml) and propidium iodide (PI, 54 µM) and incubated for 30 min at 37°C.

#### Animal experiments

All experiments with patient derived xenografts were conducted according to the institutional animal protocols and the national laws and regulations at the Experimental Pharmacology & Oncology Berlin-Buch GmbH. NOD.Cg-*Prkdc^scid^ Il2rg^tm1Sug^*/JicTac mice (Taconic) mice (female, 6-10 weeks old) were used to perform all patient-derived xenograft experiments. Prior to the experiments, patient tumors were serially transplanted in mice at least three times. Caliper measurement was used to monitor tumor growth. Tumor volume was calculated with the formula (length x width^2^) / 2. Mice were sacrificed with cervical dislocation when tumor size exceeded 1,500 mm³. LY668 was dissolved in 20% 2-Hydroxypropyl-β-Cyclodextrin in 25 mM Phosphate Buffer, pH 2. PXG-10798 was dissolved in 1% (w/v) Methyl Cellulose 400 in water, solution was vortex for 10 min and sonicated for 10 min before dosing. Mice were treated with 50 mg/kg of LY668 (p.o., daily), 0.4 mg/kg PXG-10798 (p.o., every 3^rd^ day), the combination or vehicle control for three weeks.

For pharmacokinetic studies mice were treated once with 1 mg/kg PXG-10798. Tumor tissue was collected after different time points and samples were prepared for immunoblots to evaluate degradation of CCNK.

#### Quantification and statistical analysis FASTQ file generation

Before high-throughput sequencing, the quality, quantity and size of the PCR-amplified DNA fragments of the prepared libraries were determined with a Fragment Analyzer (Thermo Fisher Scientific). All sequencing libraries were subjected to Illumina NextSeq 500 sequencing according to the manufacturer’s instructions. After base calling with Illumina’s FASTQ Generation software (v1.0.0, NextSeq 500 Sequencing), high quality PF-clusters were selected for further analysis and sequencing quality was ascertained with FastQC (v0.11.09; available online at: http://www.bioinformatics.babraham.ac.uk/projects/fastqc/).

#### ChIP Rx data processing

ChIP-Rx samples were mapped separately to the human hg19 and to the murine mm10 genome using Bowtie2 (v2.3.5.1 ^78^) using the preset parameter “very-sensitive-local”. Further, ChIP samples were normalized to the number of mapped reads in the smallest samples. Human ChIP-seq samples were either normalized relative to the spiked-in mouse reads (ChIP-Rx), or to the same number of human reads (read-normalized samples). The normalized bam files were sorted using SAMtools v1.9 ^79^ and converted to bedgraphs using bedtools genomecov (v2.30.0 ^80^). Integrated Genome Browser ^81^ was used to visualize these density files. Metagene and density plots were generated with ngs.plot.r v2.63-7 ^82^ or DeepTools v3.5.1 ^83^ using a bin size of 10 bp. *p* values were calculated by comparing the mean of the reads count via unpaired *t*-test for the specified genomic position showed in the average density plots. Plots labeled with “all genes” refer to the 57,773 genes annotated for hg19/GRCh37.p13 by Ensembl v75 (Feb 2014), all expressed genes gene set consists of 14,704 genes.

#### Total RNA sequencing data processing

Quality control of raw reads was assessed using FastQC. Reads were aligned to the human reference genome (GRCh38) using STAR (v2.7.10a) following ENCODE paired-end RNA-sequencing recommendations (ENCODE rna-seq-pipeline v0.1.0). Transcript abundance was estimated using RSEM (1.3.3) and subsequently used for differential expression analysis using DESeq2 (v1.38.3) in R. Transcripts with less than 12 counts across all nine samples were removed. Gene set enrichment analysis (GSEA) was conducted using clusterProfiler and the Molecular Signatures Database (MSigDB) v6.0 hallmark gene sets.Differential splicing analysis was performed running rMATS v.4.1. in Docker (rMATS turbo 0.1) ^41^.

#### seCLIP data processing

For seCLIP experiments, the analysis was performed according to the Skipper workflow described in ^84^. Briefly, forward read (i5) fastq files were normalized to sequencing depth using seqtk. Per condition, the same SMI subsampled using different random seeds was provided. For samples spiked with DNA, reads mapping to the DNA spike-in were identified using STAR ^85^. Accordingly, fastq files of CLIP samples were further normalized according to the number of reads mapping to the spike-in in each sample. This strategy was applied to seCLIPs shown in Figure 4. Multiplexed eCLIP samples, shown in Figure 6, were demultiplexed using UMItools ^86^ and used without further normalization. Subsequently, samples were trimmed utilizing skewer ^87^ and extracts unimolecular identifiers (UMIs) using fastp ^88^ and aligned to the human genome (GRCh38) using STAR ^85^.^85^. PCR duplicates were removed by using UMItools ^86^. To identify genomic regions with enriched signal in the CLIP over the SMI input, the automated Skipper analysis pipeline was used. Therefore, GENCODE version 38 gene annotations were obtained from gencodegenes.org. Transcripts were filtered based on total RNA-seq data in K562 cells with a minimum expression level of one transcript per million. A custom parse_gff.R script ^84^ was used to define RNA feature and transcript types (CDS_START ‘Start codon’, CDS_STOP ‘Stop codon’, CDS ‘CDS’ (protein coding genes), UTR5 ‘5’ UTR’, UTR3 ‘3’ UTR’, EXON_SMALL ‘ncRNA’ (including snoRNA, snRNA, scaRNA, scRNA, miRNA, vault_RNA, rRNA, rRNA_pseudogene, Mt_rRNA, Mt_tRNA, ribozyme, sRNA, misc_RNA), INTRON ‘Intron’, EXON_LNCRNA ‘lncRNA’ (including lncRNA, processed_transcript), EXON_MRNA ‘Other’ (including Immunoglobulin (Ig) variable chain and T-cell receptor (TcR) genes, retaind_intron, non_stop_decay, protein_cidubg_LoF, nonsense_mediated_decay, TEC), PSEUDO ‘Pseudogene’ (including inactivated immunoglobulin genes, pseudogene, polymorphic_pseudogene, translated_pseudogene, translated_unprocessed_pseudogene, pseudogene, transcribed_unprocessed_pseudogene, transcribed_processed_pseudogene, transcribed_unitary_pseudogene, unitary_pseudogene, processed_pseudogene, unprocessed_pseudogene), PRIMIRNA ‘pri-miRNA’, SS3_ADJ ‘3’ splice site adj. (100 bp intronic region upstream of the 3’ splice site), SS3_ADJ ‘5’ splice site adj. (100 bp intronic region downstream of the 5’ splice site), SSB_ADJ ’Branch point adj. (100 bp intronic region centered around the branch point), SS3_ADJ ‘5’ splice site prox. (500 bp intronic region downstream of the 5’ splice site), SS3_ADJ ‘3’ splice site prox. (500 bp intronic region upstream of the 3’ splice site), SSB_ADJ ’Branch point prox. (500 bp intronic region centered around the branch point). Each feature was tiled into evenly spaced windows of maximum 100 nucleotides in length and then tested for enrichment in immunoprecipitated over SMI samples using a beta-binomial distribution that accounts for overdispersion in read counts and GC bias. For Aurora-A seCLIP experiments the statistical thresholds were customized. Within individual replicates, windows that passed a *p* value threshold *p*<0.05 were considered enriched windows. Of these, windows that were identified in both replicates and passed a 20% FDR in at least one of the replicates were designated as reproducibly enriched windows. For the Aurora-A eCLIP in SH-EP MYCN^wt/3A^ cells 3 replicates were merged and within individual replicates, windows that passed a *p* value threshold *p*<0.05 were considered enriched windows. Of these, windows that were identified as enriched in at least two of the three replicates and passed a 20% FDR in at least one of the replicates were designated as reproducibly enriched windows. Windows that were called in more than 17% of all K562 and HepG2 eCLIP samples were blacklisted and removed. Reproducible windows were fine mapped by centering them around the local maximum of binding enrichment per window and adjusted to a fixed size of 75 nt.

To visualize similarities in binding characteristics to different RNA transcript types in a t-SNE plot, performed Aurora-A seCLIP data was pooled with ENCODE reference RBP eCLIP data, and plotted together in a two-dimensional space using the Rtsne package (https://lvdmaaten.github.io/tsne/). Several random seeds were tested to confirm a reproducible association of the queried protein with a certain feature type. Enrichment odds ratios (ORs), represented by the enrichment of observed CLIP enrichment over a baseline CLIP enrichment per RNA feature type, were calculated based on the call_enriched_windows.R script ^84^. For metagenes and heatmaps, log_2_FC of CLIPs over inputs were determined based on deduplicated bam files using bamCompare and plotted using deeptools ^89^ using the –sortUsingSamples and --averageTypeSummaryPlot mean options. Intersections between different sets of reproducible enriched windows were generated and visualized using intervene ^90^. Publicly available ENCODE eCLIP and control datasets were downloaded and log_2_FCs of CLIP over SMIs were calculated using bamCompare and plotted over called windows of interest using deeptools ^89^.

#### Mass spectrometry data analysis

MS data was analysed in Proteome Discoverer v. 2.5.0.400 using Mascot search engine v. 2.8 and UniProt Mus Musculus reviewed database (20180424; 53,102 sequences). Phospho STY and Oxidation (M) were allowed as dynamic modifications, while TMT label (N-term and K) and carbamidomethylation (C) were set as static modifications. Non-identified MS spectra was then passed through a second search using the Sequest search engine. Peptides with modification site probability <75% were filtered out. A correction for the isotopic impurity of TMT tags was applied according to the manufacturer’s correction factors for TMT batch #VL313890. Ion co-isolation threshold was set to 50% and only spectra with average S/N ratio >10 were considered for quantification. Protein abundances were normalized based on total protein amount and scaled on controls average using Proteome Discoverer’s inbuilt algorithms. Protein ratios were calculated based on the median values from all three bioreplicates of each condition. Hypothesis testing was done using ANOVA based on the abundances of individual proteins or peptides. Data from the phosphoenriched fractions was processed separately from the data from the total protein fractions. Further statistical analysis was performed in Perseus software (v. 1.6.14.0). First, filter was applied to exclude any peptides with more than 1 missing valid value in any group. Differentially phosphorylated sites were pairwise calculated by two-sided t-test with permutation-based FDR, using the default parameters. Significance was considered when *p value* =< 0.05 and log_2_(Fold Change) >= [1]. Volcano plots were generated via ggplot2 ^91^ R version 3.4.4 for macOS High Sierra (R Core Team, 2013).

#### Cell cycle immunofluorescence analysis

Cell cycle phases were determined based on Hoechst and EdU intensity. Hoechst sum staining per nucleus was used to determine the local minimum between two local maxima of cell populations representing 2n and 4n DNA content and was defined as cutoff between G1 and G2 phase. When no clear distinction between G1 and G2 phase could be made, cells were only defined as EdU^-^ or EdU^+^ (Figure S4B). EdU intensity was used to define cells as EdU^-^ (G1 and G2 phase) or EdU^+^ (S phase). Respective cutoffs were calculated according to the control condition (DMSO) of each experiment.

#### Cell viability and western blot quantification

For the quantification of live cell imaging the basic analyzer tool of the Incucyte software (v2021C) was used to quantify the confluence via phase objects. The obtained values (Mean ± SD) were visualized as percentage over time using GraphPad Prism (v10.0.0). Endpoint confluency (Mean ± SD) was visualized as bar graphs and *p* values were calculated using one-way ANOVA.

To quantify pCDK12 level (Figure 1I, Figure S1J) the immunoblot images were loaded into ImageJ (1.53t) and relative band intensities were determined and normalized to the DMSO control of each individual immunoblot to obtain relative pCDK12 intensities between 0-100.

To quantify Aurora-A levels in the sucrose gradient centrifugation (Figure 4D) band intensities were analyzed with Image Studio Lite and the signal of each fraction was normalized to total signal of all fractions.

#### Statistical analysis

*p* values for Figure 1D, 1G, 3B, 3F, 3H and S7C were calculated with one-way ANOVA using GraphPad Prism (v10.0.0). *p* values for Figures 1I, 4D, 5F and S1J were calculated with an unpaired two-sided t-test using GraphPad Prism (v10.0.0). *p* values for Figures 2F, 3C, 4A, 6B, 7C and S5A were calculated with Wilcoxon rank sum test using R (v4.2.2, r-project.org).

