## Supplemental Figures for "The MYCN/Aurora-A complex is a cyclin activating kinase for CDK12"

Figure S1, Müller et al.

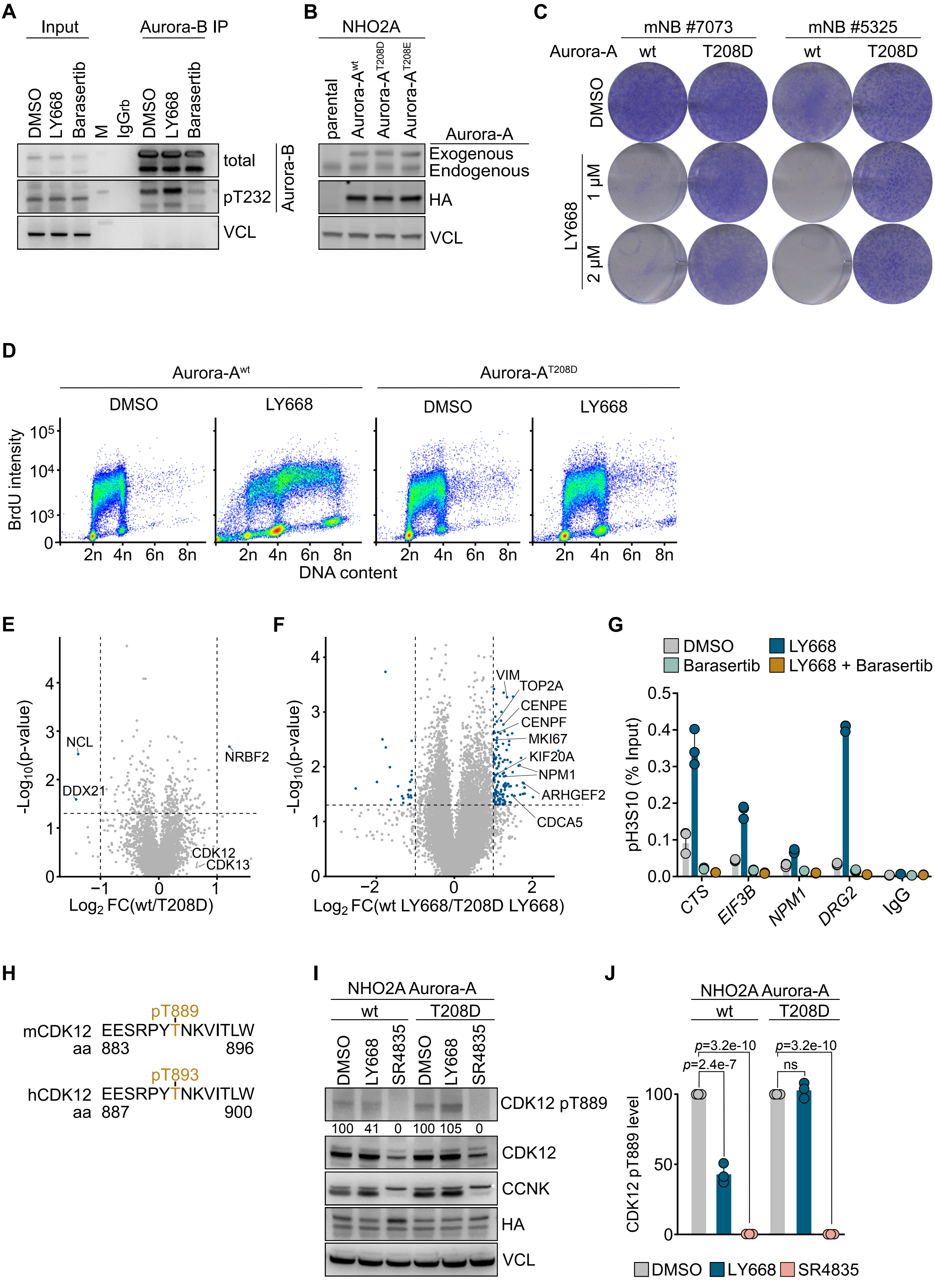

**Figure S1: Related to Figure 1.**

**A.** Immunoblots of anti-Aurora-B IPs from IMR-5 cells treated for 24 h with LY668 (1 µM), Barasertib (1 µM) or DMSO as control. The input corresponds to 1% of the amount used for the precipitation. Non-specific IgG and VCL were used as controls ($n$ = 3, M = molecular weight marker).

**B.** Immunoblots of indicated proteins in parental NHO2A and NHO2A expressing Aurora-$A^{wt}$, Aurora-$A^{T208D}$ or Aurora-$A^{T208E}$ cells. VCL was used as a loading control ($n$ = 3).

**C.** Clonogenic assay in murine neuroblastoma cells established from the tumor of two different TH-MYCN mice expressing Aurora-$A^{wt}$ or Aurora-$A^{T208D}$ treated with LY668 or DMSO as a control ($n$ = 3).

**D.** BrdU/PI flow cytometry profiles of NHO2A cells expressing Aurora-$A^{wt}$ or Aurora-$A^{T208D}$ treated with LY668 or DMSO as a control ($n$ = 3).

**E.** Volcano plot of quantitative phospho-proteomics data comparing the effect in control conditions on NHO2A cells expressing either Aurora-$A^{wt}$ or Aurora-$A^{T208D}$. The x axis displays phospho-peptide $log_2$Fold change (FC) between cells expressing either Aurora-$A^{wt}$ or Aurora-$A^{T208D}$. The y axis shows statistical significance ($n$ = 3).

**F.** Volcano plot of quantitative phospho-proteomics also shown in Figure 1E. Marked are sites enriched in NHO2A expressing Aurora-$A^{wt}$ upon LY668 treatment.

**G.** pH3S10 ChIP-qPCR at indicated *loci* in IMR-5 cells after treatment as indicated for 6 h. IgG was used as negative control. Technical triplicates and the mean of one representative experiment are shown ($n$ = 3).

**H.** Scheme showing the homology between the murine and the human T-loop sites of CDK12. Highlighted are the phospho-sites.

**I.** Immunoblots of indicated proteins in NHO2A cells expressing Aurora-$A^{wt}$ or Aurora-$A^{T208D}$ upon treatment with LY668 (1 µM) or SR4835 (200 nM) for 8 h. Asterisks indicate unspecific bands. VCL was used as a loading control ($n$ = 3).

**J.** Quantification of CDK12 pT889 levels in replicate immunoblots described in H. Data are shown as mean ± SD. $p$ values are calculated using an unpaired two-sided t-test ($n$ = 3).

**Figure S2, Müller et al.**

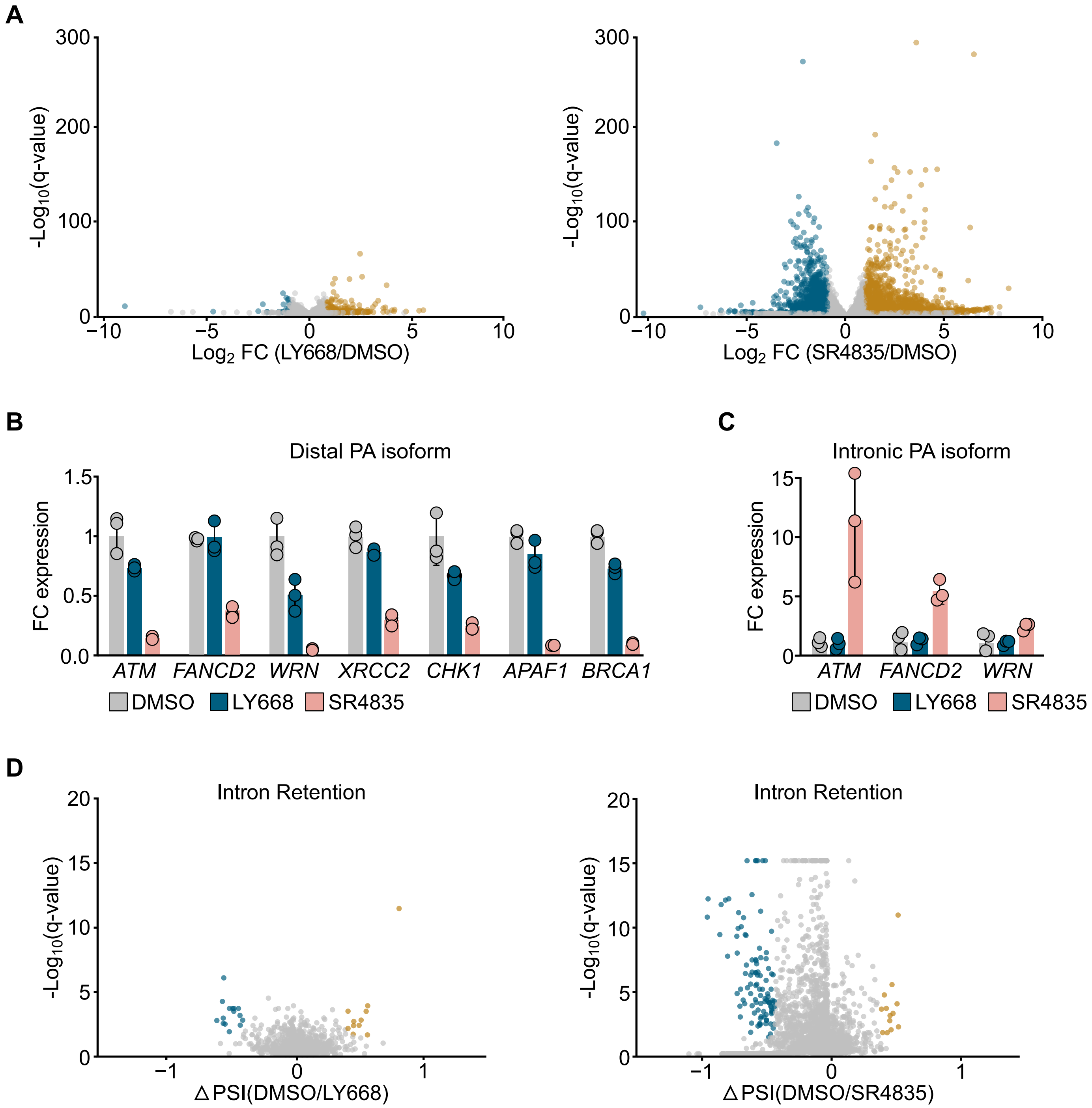

**Figure S2: Related to Figure 2.**

**A.** Volcano plot of total RNA sequencing data showing differences in gene expression after treatment for 24 h with LY668 (1 µM, left) or SR4835 (50 nM, right) compared to DMSO as a control. Colored dots indicate significantly up- or downregulated genes. The x axis shows the $log_2FC$ between treated and control cells, and the y axis depicts statistical significance (*n* = 3).

**B.** Relative expression of distal polyadenylation (PA) isoforms of indicated genes in IMR-5 cells treated with LY668 (1 µM), SR4835 (100 nM) or DMSO as control for 8 h (*n* = 3).

**C.** Relative expression of intronic PA isoforms of indicated genes in IMR-5 cells treated as described in B (*n* = 3).

**D.** Splicing analysis using rMATS measuring intron retention in IMR-5 cells treated with LY668 (1 µM, left), SR4835 (50 nM, right) or DMSO for 24 h. Colored dots indicate significantly affected genes. The x axis shows the delta percent spliced in (∆PSI) between treated and control cells, and the y axis depicts statistical significance (*n* = 3).

**Figure S3, Müller et al.**

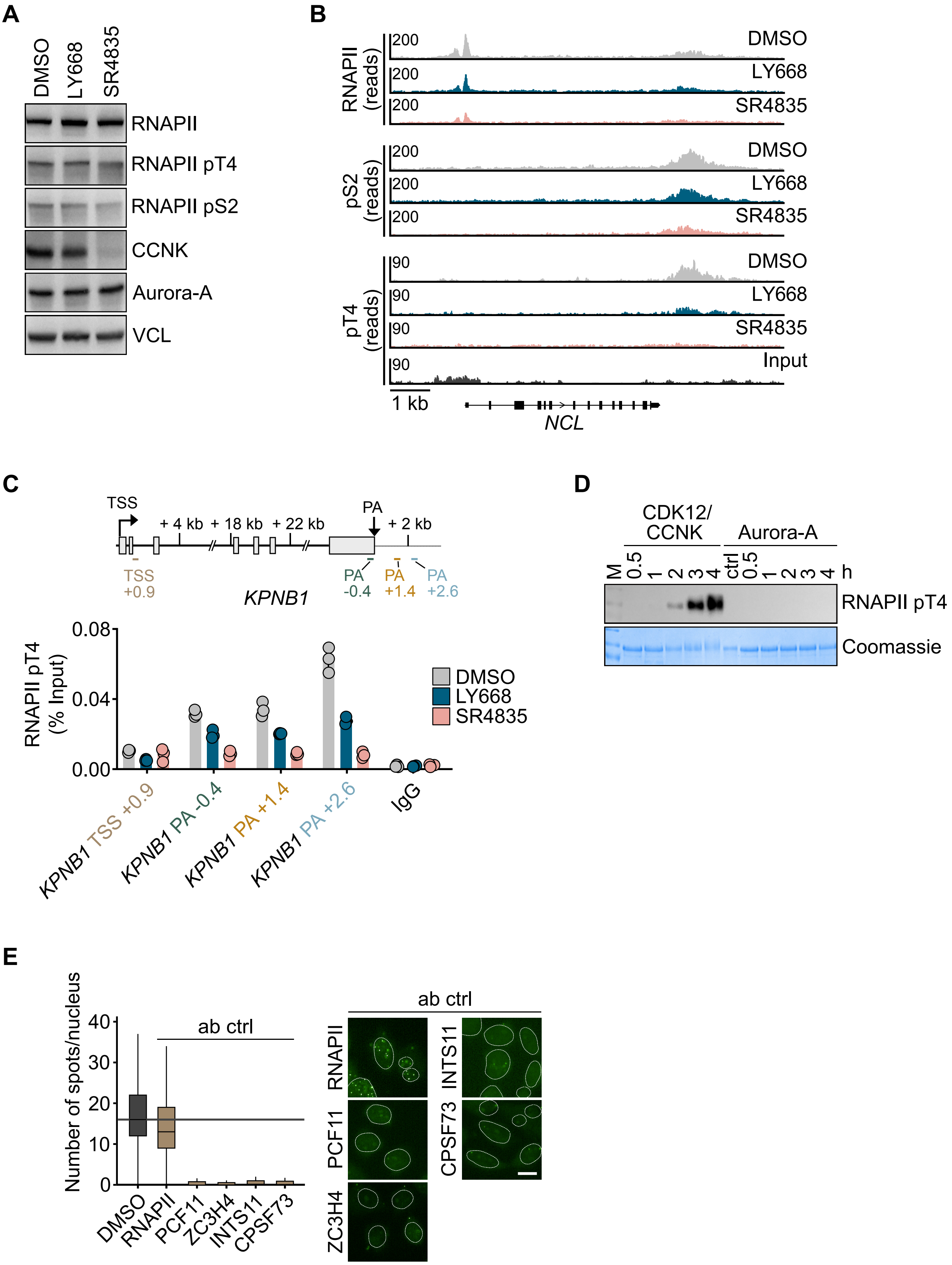

**Figure S3: Related to Figure 2.**

**A.** Immunoblot of indicated proteins in IMR-5 cells treated for 24 h with LY668 (1 µM), SR4835 (50 nM) or DMSO as a control. VCL shown as loading control (*n* = 3).

**B.** Genome browser tracks reporting the chromatin occupancy of RNAPII (total, pS2 or pT4) at the *NCL* gene in IMR-5 cells treated as described in A (*n* = 2 for total RNAPII and pS2; *n* = 1 for pT4).

**C.** RNAPII pT4 ChIP-qPCR at the *KPNB1* locus in IMR-5 cells after treatment as indicated for 24 h. IgG was used as negative control. Primer locations are shown in the schematic above. Shown are technical triplicates and the mean of one representative experiment (*n* = 3).

**D.** *In vitro* kinase activity experiment with subsequent immunoblotting for RNAPII pT4. Coomassie staining is shown as loading control (*n* = 1).

**E.** Quantification and representative images of single antibody controls (ab ctrl) for PLAs shown in Figure 2F, G. DMSO refers to PLA signal with the lowest measured spots per nucleus. Scale bar indicates 10 µm.

**Figure S4, Müller et al.**

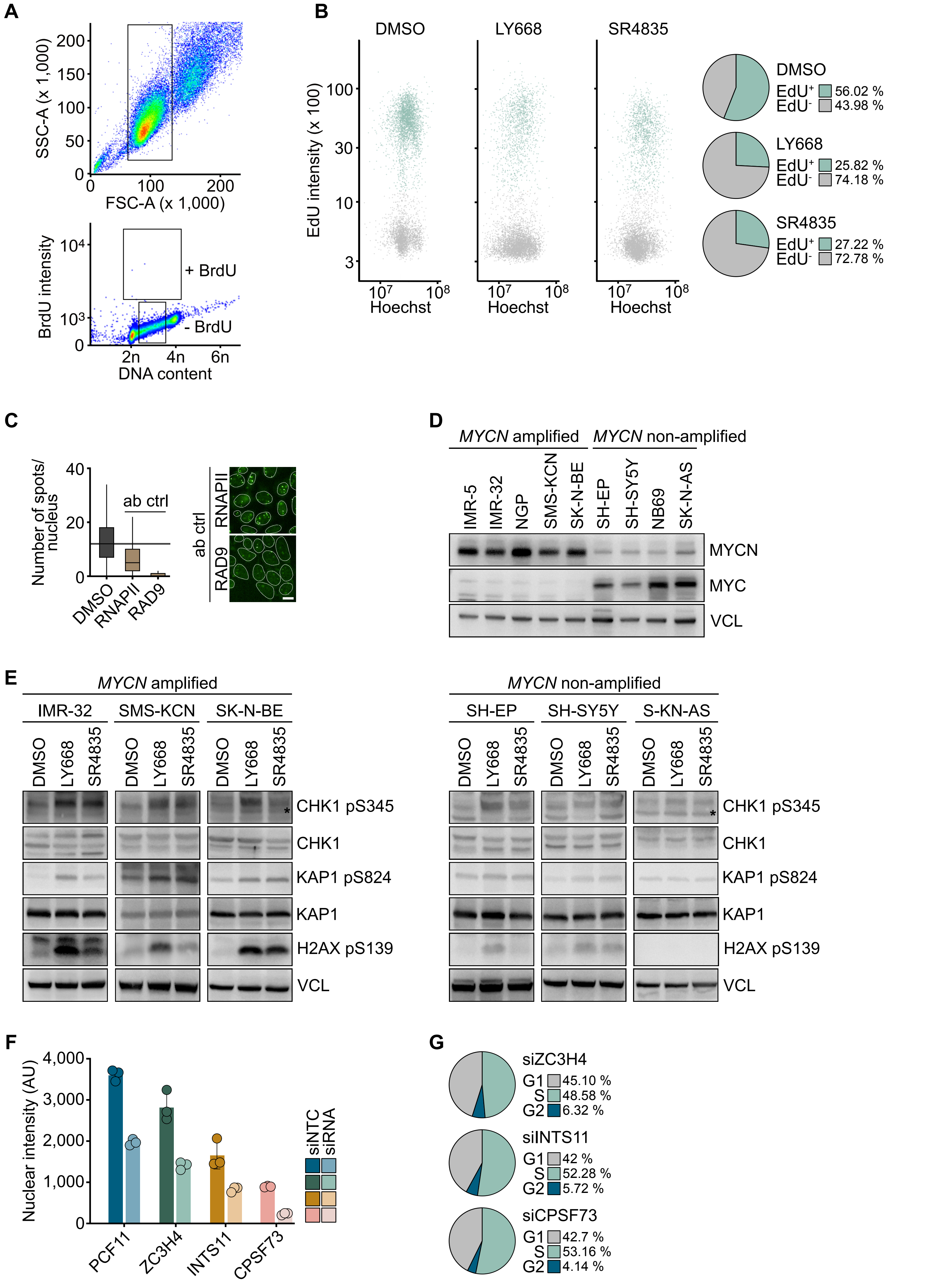

**Figure S4: Related to Figure 3.**

**A.** Gating strategy for viable IMR-5 cells (top) and for BrdU/PI analysis (bottom) shown in Figure 3A, B on a control sample without BrdU staining (*n* = 3).

**B.** (Left) Immunofluorescence-based cell cycle analysis of IMR-5 cells treated as indicated for 24 h. The x axis displays the Hoechst intensity, and the y axis depicts EdU incorporation. (Right) Quantification of one representative experiment (*n* = 3).

**C.** Quantification and representative images of single antibody controls (ab ctrl) of PLAs shown in Figure 3C, D. Scale bar indicates 10 µm (*n* = 3).

**D.** Immunoblot of MYCN and MYC level in a panel of neuroblastoma cell lines. VCL was used as a loading control (*n* = 2).

**E.** Immunoblots of indicated proteins in *MYCN* amplified (left) and *MYCN* non-amplified (right) neuroblastoma cells upon treatment with LY668 (1 µM), SR4835 (50 nM) or DMSO as a control for 24 h. Asterisks indicate unspecific bands. VCL was used as a loading control (*n* = 2).

**F.** Nuclear intensity of indicated proteins after transfection with corresponding siRNAs compared to siNTC. Shown are technical triplicates and the mean of one representative experiment (*n* = 3).

**G.** Representative cell cycle distribution of IMR-5 cells determined by immunofluorescence upon transfection with siZC3H4, siINTS11 or siCPSF73 (*n* = 3).

**Figure S5, Müller et al.**

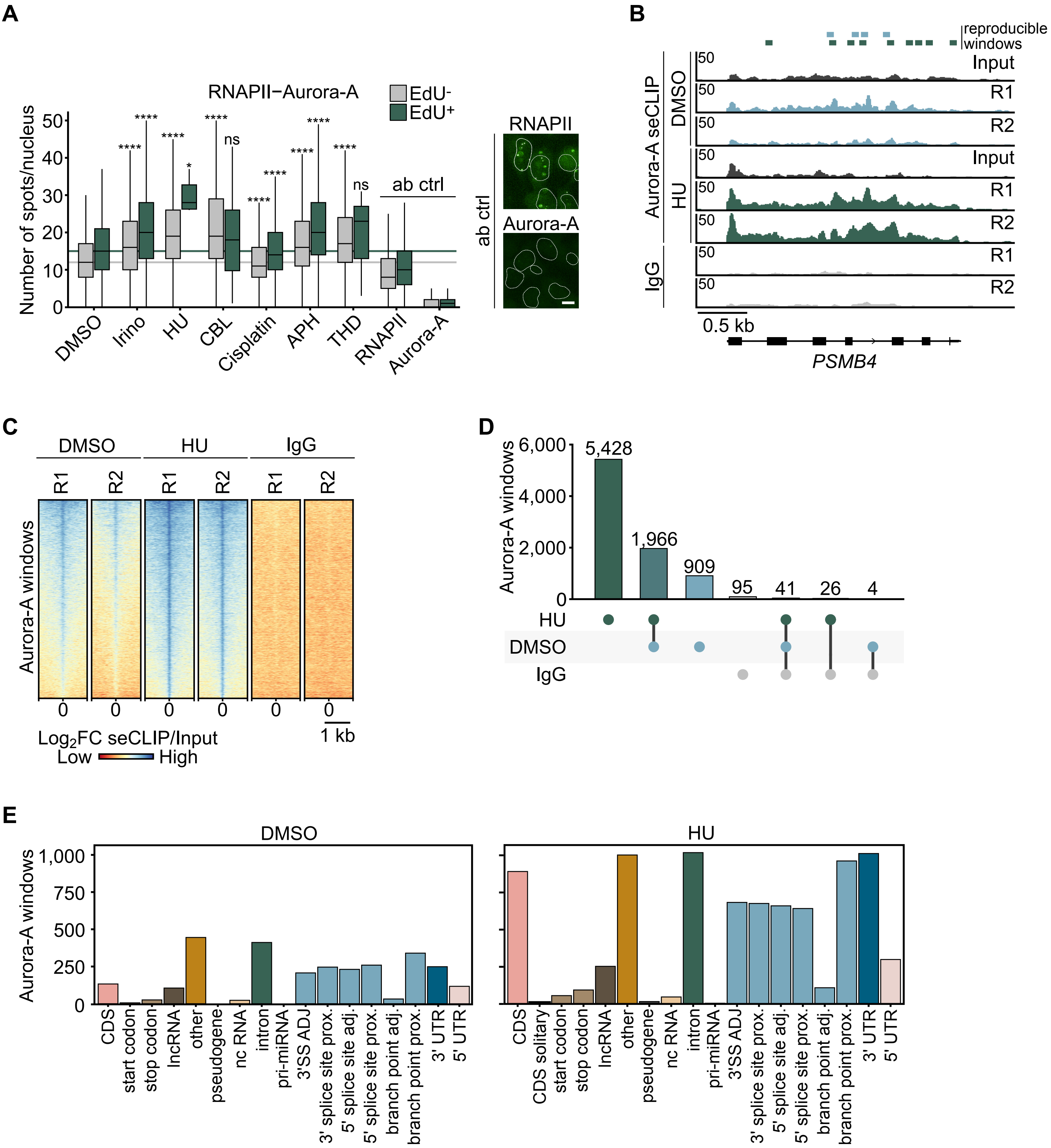

**Figure S5: Related to Figure 4.**

**A.** (Left) Boxplot of single-cell analysis of nuclear PLA foci between RNAPII and Aurora-A in IMR-5 cells treated as indicated. Conditions are stratified for EdU incorporation. *p* values were calculated comparing the PLA signal of all analyzed cells using Wilcoxon rank sum test compared to corresponding control (*$p$ value<0.5; ****$p$ value<1.0e-15; $n$ = 2). (Right) Representative images of single antibody controls (ab ctrl). Scale bar indicates 10 µm.

**B.** Genome browser tracks of Aurora-A seCLIPs at the *PSMB4* locus. IMR-5 cells were treated with HU or DMSO as a control. Input and IgG are shown as control. Bars on top show reproducible enriched windows ($n$ = 2, R = replicate).

**C.** Heatmaps showing the $\log_2$FC of Aurora-A seCLIPs over the respective size-matched input in depicted conditions, sorted for merged Aurora-A DMSO/HU windows.

**D.** UpSet plot of Aurora-A seCLIP windows comparing enriched windows in HU, DMSO and IgG as a control.

**E.** Bar chart showing classification of RNA species in enriched windows of Aurora-A seCLIPs.

**Figure S6, Müller et al.**

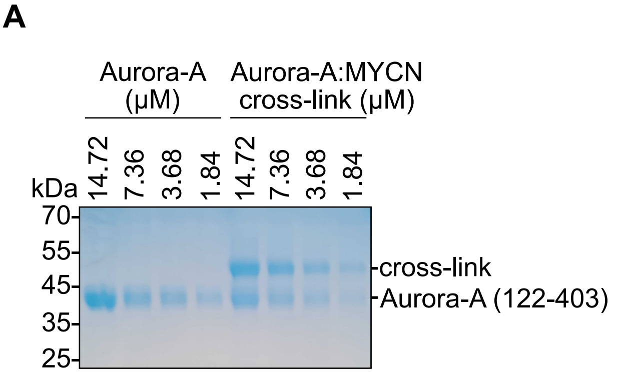


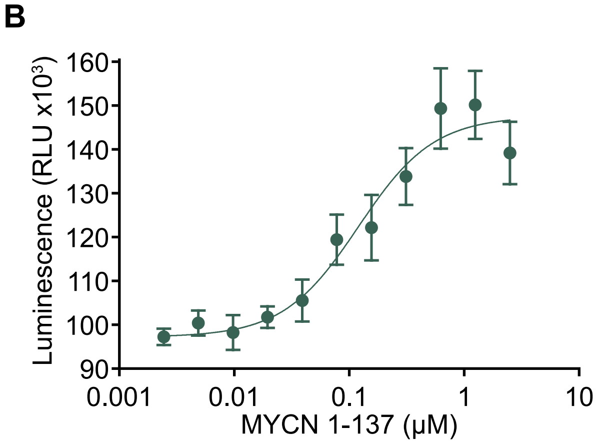


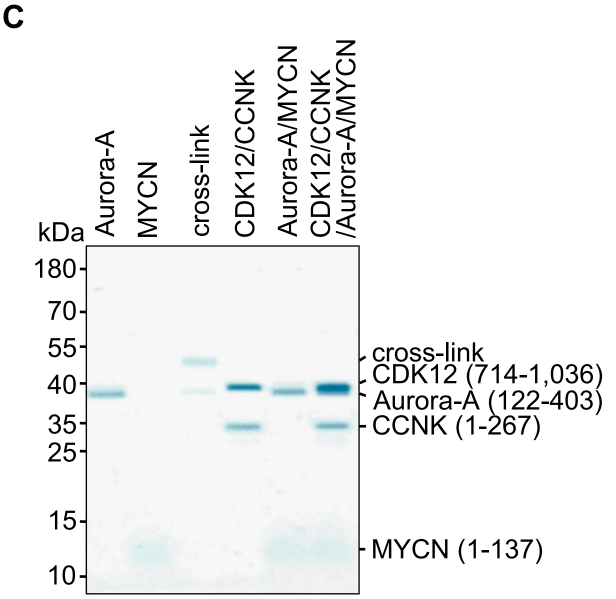

**Figure S6: Related to Figure 5.**

**A.** Coomassie staining of recombinant Aurora-A or Aurora-A:MYCN cross-link. Note that Aurora-A:MYCN cross-link contains free Aurora-A (lower band) (*n* = 2).

**B.** *In vitro* ADP-Glo kinase assay reporting activity of initially unphosphorylated Aurora-A aa 1-403 with a 2-fold serial dilution of MYCN (1-137, 2.5 μM – 2.4 nM). The EC50 for Aurora-A activation by MYCN 1-137 was 120 ± 22 nM. Data is shown as mean ± SD (n = 3).

**C.** Coomassie staining of indicated recombinant proteins used for kinase assay shown in Figure 5E. Note that CDK12 and Aurora-A migrate very close to each other (n = 2).

**Figure S7, Müller et al.**

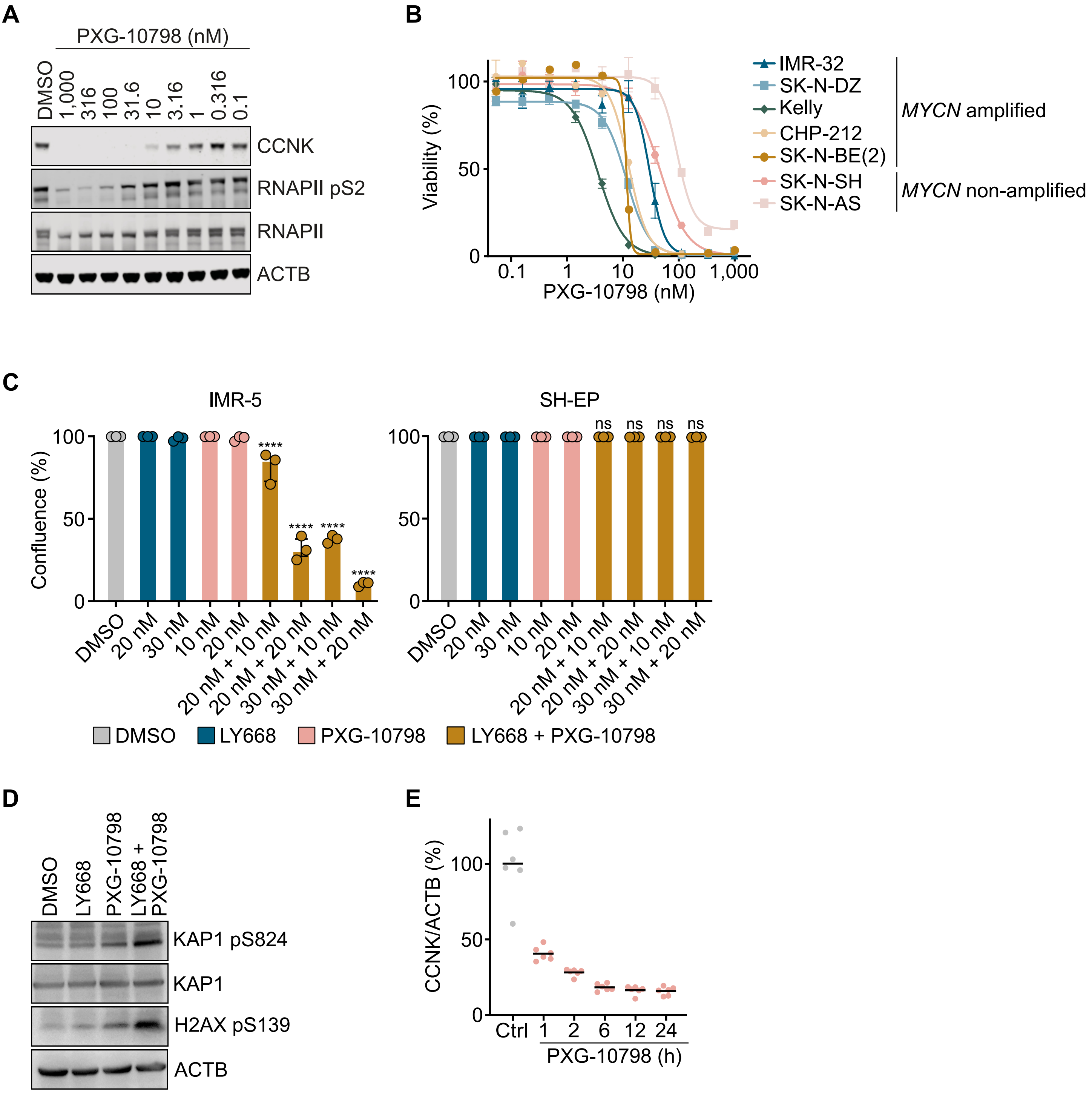

**Figure S7: Related to Figure 7.**

**A.** Immunoblot of indicated proteins in BT474 cells treated with indicated concentrations of PXG-10798 (*n* = 3).

**B.** Cell viability of *MYCN* amplified (left) and *MYCN* non-amplified (right) neuroblastoma cells upon treatment with increasing concentrations of PXG-10798. Shown is the mean of three technical replicates.

**C.** Endpoint confluence from growth curves of IMR-5 or SH-EP cells treated with different concentration of LY668, PXG-10798 or the combination as indicated. Data shown as mean ± SD. *p* values were calculated using an one-way ANOVA (*****p* value<0.0001; *n* = 3)

**D.** Immunoblots of indicated proteins in IMR-5 cells upon treatment with LY668 (30 nM), PXG-10798 (15 nM), the combination or DMSO as a control for 48 h. ACTB was used as a loading control (*n* = 3).

**E.** PD profile depicting the ratio of CCNK/ACTB after one time treatment with PXG-10798 at 1 mg/kg measured after indicated time points (N = 6 mice per group).

**Table S1** Table providing significant phosphorylation sites ($log_2FC<-1$, $p$ value<0.05) of quantitative phospho-proteomics data comparing the effect of LY668 on NHO2A cells expressing either Aurora-$A^{wt}$ or Aurora-$A^{T208D}$.

| Gene | Modifications in Master Proteins | Log2FC(WT LY668 8h/T206D LY668 8h) | -log10(p-value) |
|---|---|---|---|
| Cdk12 | [T889(100)] | -1,7676182 | 3,73406142 |
| Cdk13 | [T871(100)] | -1,8437417 | 2,50102412 |
| Rpl7l1 | [S230(100)] | -1,0936157 | 2,47325898 |
| Rbmxl1 | [S212(99.8)] | -1,757737 | 2,35297669 |
| Slc39a10 | [T542(100)] | -1,6525233 | 1,99305296 |
| Ltv1 | [S347(100);S350(99);T351(99);S353(100)] | -1,0889632 | 1,92195742 |
| Sun2 | [S120(99.6);S/T] | -1,0976257 | 1,85023179 |
| Hnrnpul2 | [S166(100)] | -1,0923367 | 1,76537832 |
| Tmpo | [S384(100)] | -1,1317056 | 1,7250292 |
| Ddx23 | [S106(99.4);S108(100)] | -1,9853854 | 1,72227909 |
| Smarca5 | [S65(100)] | -1,4439359 | 1,64428411 |
| Dkc1 | [S508(100)] | -2,5304413 | 1,60393688 |
| Srsf1 | N/A | -1,1240612 | 1,57699157 |
| Palm | [T145(100)] | -1,1654806 | 1,57272809 |
| Ralbp1 | [S29(99.6)] | -1,1886457 | 1,49074315 |
| Ctnnd1 | [T788(100)];[T815(100)] | -1,0040545 | 1,47741306 |
| Ddx54 | [S774(100)] | -1,118205 | 1,47637156 |
| Srrm1 | [S723(100);S725(100);S731(100)] | -1,3413662 | 1,46476536 |
| Mtres1 | [S106(100)] | -1,1720235 | 1,44363694 |
| Ccnl1 | [S380(100)] | -1,1052738 | 1,43785044 |
| Pinx1 | [T228(100);S233(100)] | -1,3105415 | 1,42032733 |
| Prkcd | [T390(99.6)] | -1,6278526 | 1,39995254 |
| Vim | [S339(100)] | -1,3100343 | 1,34124495 |
| Znf280c | [S528(100)] | -1,1373311 | 1,31871475 |
